# A cortical gradient of distance to criticality governs large-scale resting-state fMRI dynamics

**DOI:** 10.64898/2026.05.21.726898

**Authors:** D. Yellin, E. Simony, R. Malach, O. Shriki

## Abstract

The cortex generates a remarkable diversity of regional dynamics despite relying on broadly shared principles of recurrent organization. Here, we identify distance to criticality (DTC) as a unifying axis along which these dynamics are spatially organized. Analyzing resting-state fMRI BOLD signals and leveraging simple network models of randomly connected recurrent units, we show that DTC robustly explains key dynamical features, in particular, local power spectra and functional connectivity, across the full set of 360 cortical areas. Our analysis shows that a rank-order distribution of DTC values is highly conserved across subjects. Moreover, the empirical analysis of cortical slow dynamics and its fitted network simulations demonstrate similar power-laws across hierarchies of the cortical sheet. These results suggest that recurrent neuronal networks, operating close to criticality, can generate a remarkably rich dynamical repertoire which fit the entire range of experimentally observed cortical slow-timescale dynamics. Our findings underscore the importance of DTC as a powerful, fundamental generator underlying the spectrum of diverse cortical dynamics.

**Highlights:**

- Spontaneous (resting-state) activity in the human cortex is shown to be organized along a conserved spatial gradient of distance from criticality (DTC), with regions exhibiting a stable cross-individual rank order along this axis.
- Multi-subject fMRI data of regional power spectra and functional connectivity can be fitted with a single parameter simulation model based on DTC.
- Quantitative estimation of the DTC across cortical regions can be achieved using a simple sparse recurrent neural network model.
- The model fits the power spectra of low frequency fluctuations and the distribution of functional connectivity.
- Shape collapse analysis of the power spectrum demonstrates a universal profile across the resting cortex depending only on the DTC.

## Introduction

One of the most striking puzzles of the cerebral cortex is its capacity to underlie highly diverse dynamics reflecting a large range of characteristic timescales. These dynamics are organized along different hierarchies dissected into interrelated networks. Yet this extraordinary diversity is grounded in a remarkably uniform anatomical structure, consisting of six layers, where the micro-distinctions between areas are rather subtle (Amir et al., 1993; Carlo & Stevens, 2013; V. Mountcastle, 1997; V. B. Mountcastle, 1957).

A noteworthy and significant aspect of these dynamics is reflected in their wide-spread nature across the cortical mantle, in the form of spontaneous fluctuations. In fact, these fluctuations have been found in each and every cortical region studied (Smith et al., 2015). Termed spontaneous or resting state fluctuations, these dynamic phenomena are surprisingly slow, taking seconds to evolve and obeying a largely power-law spectral signature (He, 2014; Nir et al., 2008). Because of their ultra-slow dynamics, such fluctuations have been amply demonstrated also using non-invasive BOLD-fMRI (Biswal et al., 1995; Fox & Raichle, 2007; Nir et al., 2007; Raichle, 2011; Yellin et al., 2015). Indeed, a large body of research has demonstrated such ultra-slow, scale-free dynamics of neuronal fluctuations across the cerebral cortex (He, 2014; Rolls et al., 2021). Given the fast nature of neuronal activations, typically appearing in a fraction of a second (Allison, 1999; Fisch et al., 2009; Thorpe et al., 1996), the ultra-slow nature of these fluctuations poses a major puzzle. Importantly, despite their wide-spread nature, these fluctuations differ significantly in their specific dynamic signature across cortical sites (Broday-Dvir & Malach, 2021; Honey et al., 2012; Kucyi et al., 2018). This has led to a number of theoretical studies proposing a mechanism that will explain the emergence of such slow cortical dynamics.

Besides the ultra-slow dynamics, another particularly informative aspect of the spontaneous fluctuations is the fact that they emerge in correlated patterns, termed resting-state networks. These patterns are revealed, experimentally, through temporal correlation structures also termed “functional connectivity” (Biswal et al., 1995; Fox et al., 2006; Nir et al., 2008). These connectivity structures have been linked to task-related networks (Fox et al., 2006; Tommasin et al., 2017) as well as habits and prior experiences (Harmelech & Malach, 2013), and have been the focus of a large research effort by many groups. As in the case of spontaneous fluctuations, the functional connectivity across cortical sites also changes substantially, possibly reflecting the underlying difference in anatomical intrinsic connectivity (Amir et al., 1993; Keller et al., 2013; Kucyi et al., 2018).

Here, we propose that a central factor, linking this uniform structure to the diverse dynamics, stems from the unique recurrent architecture, which is a major landmark of cortical circuitry (Douglas & Martin, 2007; Morishima & Kawaguchi, 2006). An important manifestation of such recurrence, is its impact on the neuronal dynamics, allowing the system to approach a critical phase transition (Hahn et al., 2017; Ponce-Alvarez et al., 2023). We and others have recently used recurrent network simulations and successfully demonstrated the emergence of ultra-slow dynamics in such networks associated with proximity to criticality (Deco & Jirsa, 2012; Gavenas et al., 2024; Spiegler et al., 2016; Yellin et al., 2025).

Given this “critical brain hypothesis” (Beggs, 2008; Beggs & Plenz, 2003; Chialvo & Bak, 1999) which posits that the cortex operates near critical dynamics, can we quantify how close different cortical regions are to criticality, and is this proximity organized along a systematic distribution across the brain? The question addressed in this paper is whether local proximity to criticality, a fundamental dynamical property of recurrent cortical circuitry, can account for two key features of resting-state activity across the cortex: the ultraslow spontaneous fluctuations observed in regional fMRI power spectra and the spatial profile of functional connectivity. Specifically, we ask whether differences in the distance of local networks to the critical regime can explain the diversity of fluctuation dynamics and correlation structures observed across cortical regions. This issue is relevant to a recently emerging subfield of study termed ‘distance to criticality’ (DTC; Liu et al., 2023; Muñoz, 2018; O’Byrne & Jerbi, 2022; Wilting & Priesemann, 2015), which employs a range of methods to assess the proximity to criticality across distinct neuronal populations. We note that although DTC is the common and prevailing terminology in the literature, we occasionally employ the alternative term proximity to criticality (PTC; Sooter et al., 2025) to emphasize intuitive closeness to the phase transition. This terminological choice is adopted for conceptual clarity only. In this sense, DTC decreases as the system approaches the phase transition, whereas PTC increases, reflecting complementary descriptions of the same underlying notion. Nonetheless, the DTC distribution across the human cortex remains poorly understood. Here, we hypothesize that the slow and coordinated fluctuations in spontaneous cortical activity may capture a signature of near-critical dynamics, facilitating the estimation of DTC across cortical areas.

In this work we focus on instability criticality, a type of dynamical phase transition in recurrent networks that separates stable activity, where perturbations decay, from unstable dynamics in which activity grows explosively (Beggs, 2008; Palva et al., 2013). Other forms of critical transitions, such as those leading to chaotic dynamics or pattern formation, have been proposed in neural systems ( O’Byrne & Jerbi, 2022; Algom & Shriki, 2026). Near the critical boundary, the effective restoring forces that stabilize activity become weak, leading to a strong amplification of intrinsic timescales through recurrent interactions. As a result, perturbations relax slowly and correlations decay over extended timescales, giving rise to long-range temporal correlations with approximate power-law scaling, reflecting the dominance of slow collective modes near the critical boundary. This phenomenon is termed ‘critical slowing down’ (CSD) (Dotan & Shriki, 2021; Gavenas et al., 2024; Meisel et al., 2015; Shriki & Yellin, 2016; Yellin et al., 2025).

Based on simulation results from former work (Yellin et al., 2025), we observed that as the network approaches criticality - (a) the amplitude of slower fluctuations grows larger (due to CSD), and, (b) the mean correlation within the local neuronal population becomes stronger, as illustrated in Figure 1. The random recurrent network (Figure 1A) displays the spectral dynamics leading to CSD when the neural gain approaches criticality (Figure 1B). The power spectral density (PSD) assumes a power-law scaling (i.e. approaching a straight line on the PSD log-log plot illustrated in Figure 1C). The local population firing rate displays amplification of coherent ultra-slow fluctuations (demonstrated in Figure 1D) as the network operates closer to criticality.

**Figure 1.**
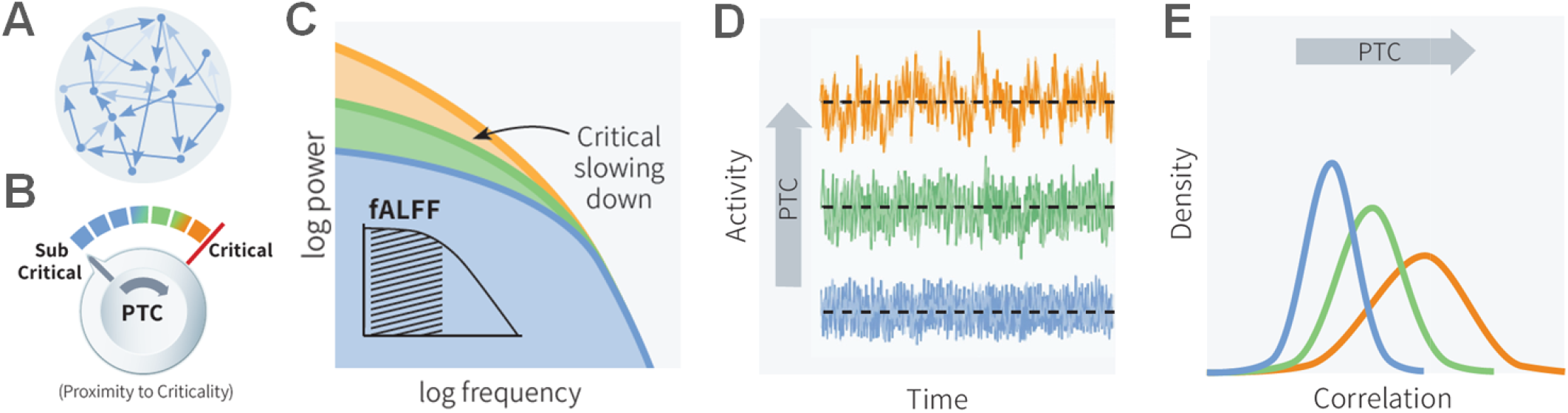
Proximity to criticality (PTC) shapes ultra-slow fluctuations and local correlations in recurrent networks. (A) Schematic of a random recurrent network model. (B) Neural gain is modulated by a control parameter (knob), which determines the network’s PTC and is color-coded consistently across panels, ranging from blue (sub-critical), through green, to orange as the system converges toward the critical point. (C) As PTC increases, the power spectral density (PSD) of network activity transitions toward scale-free behavior, exhibiting an approximate power-law scaling visible as a straight line in log– log coordinates. An inset illustrates the corresponding fractional amplitude of low-frequency fluctuations (fALFF), defined as the fraction of power in the low-frequency band of the PSD. (D) Increased PTC is accompanied by amplification of coherent ultra-slow fluctuations in the local population firing rate. (E) The same progression leads to enhanced local functional connectivity, reflected in a systematic broadening and strengthening of the distribution of pairwise population correlations. Note the increase in the amplitude of the slow fluctuations and the increase in functional connectivity as the system approaches criticality.

Figure 1E illustrates the increase in local functional connectivity (observed through the population’s distribution of pairwise correlations) as the system approaches criticality.

To explore this relation further, we analyzed a large scale dataset of resting state fluctuations at high resolution (7T) fMRI that were collected in the Human Connectome Project (HCP) (Barch et al., 2013; Glasser et al., 2013) and explored to what extent these data can be fitted by modifying a single DTC-related parameter in a simple random recurrent network simulation.

We take advantage of two readily available, highly explored metrics, used in fMRI data analysis to characterize measures of the abovementioned cortical ultra-slow fluctuations and activity correlations, namely, the fractional Amplitude of Low-Frequency Fluctuations (fALFF) and the Functional Connectivity Density (FCD), respectively. The measure of fALFF, reflecting the normalized power within low-frequency bands of the BOLD signal, was first suggested by (Zou et al., 2008) as a refinement of the earlier unnormalized measure of Amplitude of Low-Frequency Fluctuations (ALFF) (Yang et al., 2007; Yu-Feng et al., 2007) that expresses absolute power. Both ALFF and fALFF provide a rs-fMRI method for quantifying the relative magnitude of ultra-slow spontaneous brain activity. To address oversensitivity of ALFF to physiological noise, fALFF normalizes the power in the low-frequency range by the total power across a wider frequency spectrum. Thereby it was found to enhance specificity to neuronal activity (Huck et al., 2023; Zou et al., 2008), effectively suppressing nonspecific signal components, such as venous biases. Studies have utilized both ALFF and fALFF to investigate various neurological and psychiatric conditions, providing insights into the underlying neural mechanisms (Chen et al., 2022, 2022; Di et al., 2013; Jia et al., 2020).

The FCD measure, introduced by Tomasi and Volkow, quantifies fMRI voxels’ connectivity cardinality, i.e. the number of connections a given voxel has with its neighboring voxels (Tomasi & Volkow, 2010, 2011). FCD identifies local network hubs by assessing the degree of connectivity within a specified radius, providing insight into the brain’s functional architecture without the need for predefined seed regions. Building on this framework, we extended the concept of FCD to a regional scale, yielding a measure termed regional Functional Connectivity Density (rFCD). In this adaptation, the FCD is computed within each predefined cortical region of interest (ROI), such that the distribution (or histogram) of voxel-wise functional connectivity values is assessed across all voxels belonging to that region. This approach retains the core rationale of the original FCD—quantifying local connectivity density—while tailoring it to the internal organization of functionally defined cortical areas. The resulting rFCD thus reflects the intrinsic connectivity structure within each ROI, providing a region-specific index of local network dynamics and integration (Raut et al., 2020).

In this study, we treat the cortex as a collection of isolated functional regions, where each region of interest (ROI) is analyzed independently in terms of its fALFF and rFCD measures and simulated as a separate recurrent network. This simplification deliberately omits the extensive inter-regional connectivity characteristic of the human cortex, thereby focusing on the intrinsic dynamics governing spontaneous activity within individual areas. The principal advantage of this approach lies in its conceptual clarity and computational simplicity, enabling direct examination of how local network parameters shape regional fMRI-derived metrics. It is motivated by the well documented dense local connections found throughout the cortical mantle (Amir et al., 1993). However, this abstraction necessarily excludes large-scale interactions such as long-range synchronization, feedback modulation, and activity propagation that arise from long-range distributed connectivity. Consequently, while the isolated-ROI framework provides a controlled setting for identifying local determinants of fALFF and rFCD variability, it does not capture the large-scale integrative processes underlying whole-brain coordination. Interestingly, despite this oversimplification, our results show a good empirical-to-model fit of both dynamics and connectivity, emphasizing the role of local network processing in mediating the distribution of cortical DTC.

## Results

Our study involved the analysis and related simulation of high resolution resting state 7T fMRI data obtained in the HCP project (Barch et al., 2013; Glasser et al., 2013), employing the Glasser parcellation to 180 ROIs (Glasser et al., 2016). The brain imaging data included healthy young adults (22–35 years; N = 40; 27 females) who were instructed to fixate on a screen cursor during rest for a duration of 300 seconds (Methods). The analysis focused on two main aspects of local ROI fMRI data: the fluctuation dynamics and functional connectivity of the BOLD signal. To that end, we first examined fALFF (a representative metric of the slow-evolving fluctuation) and regional FCD (rFCD, a metric of the distribution of local functional connectivity) in each of the ROIs (Methods).

In order to model these two metrics, we simulated a highly simplified sparse random recurrent network, implemented as a firing rate model (Methods), founded on (Chaudhuri et al., 2018; Yellin et al., 2025). Each simulated network corresponded to a single fMRI ROI, with the activity of each voxel represented by the firing rate of a single unit. The analogous fALFF and rFCD measures were then computed directly from the network’s population activity (Methods). Importantly, since the HCP data consists of BOLD fMRI signals, which represent a low-pass filtered version of the neuronal firing-rate transients (Logothetis et al., 2001; Mukamel et al., 2005), we convolved the simulated neuronal activity with a hemodynamic function (Methods). For single-subject comparisons, the number of units in each simulated network was matched to the corresponding ROI number of voxels. However, for larger-scale (full population) analyses, network size was fixed at 400 units to allow for cross-regional cortical comparisons (Methods).

The proximity to the critical transition point between decaying and diverging dynamics can be evaluated using tools from random matrix theory, assuming a large enough network size (Methods). Specifically, the proximity to criticality was directly modulated through a control parameter, G, corresponding to the product of the network sparseness *p*, the neural gain, *γ*, and the mean synaptic strength *μ* _conn_, with the critical transition from subcritical to supercritical dynamics expected at G_crit_ = *γ* p*μ* _conn_ = 1 (Methods). Although the values of *p*, *γ*, and *μ* _conn_ that comprise G can vary to satisfy the near-criticality condition, we mainly used the neural gain (*γ*) to adjust the network’s proximity to the critical phase transition. The network sparseness (*p*) was found to directly determine the width of the correlation distribution curve and was thus adjusted as needed to fit the empirically observed rFCD. Modifying the last parameter controlling the mean synaptic strength was roughly equivalent to change of gain and was therefore fixed at *μ* _conn_ = 49.881 for all presented results. The time constant (τ) in these simulations was set to 20 ms, which is close to well-known estimates for cortical neurons’ membrane time constant (Koch et al., 1996; Softky & Koch, 1993; Yellin et al., 2025).

The capacity of the artificial network to generate a range of local dynamics and correlation profiles allowed us to examine the central question of this work - to what extent the spontaneous fluctuations generated by local (“isolated island”), simple, artificial recurrent networks can account for the diversity of dynamic profiles actually measured across the entire human cerebral cortex.

Figure 2 demonstrates a qualitative comparison between empirical and simulated results. Figure 2A illustrates single-subject examples of different BOLD fMRI resting state fluctuation profiles obtained from three different cortical regions (premotor, high-order visual and default-mode, low-to-high fALFF respectively), and Figure 2B presents the qualitative fits of the fluctuations generated by artificial network simulations at varying proximity to criticality.

**Figure 2.**
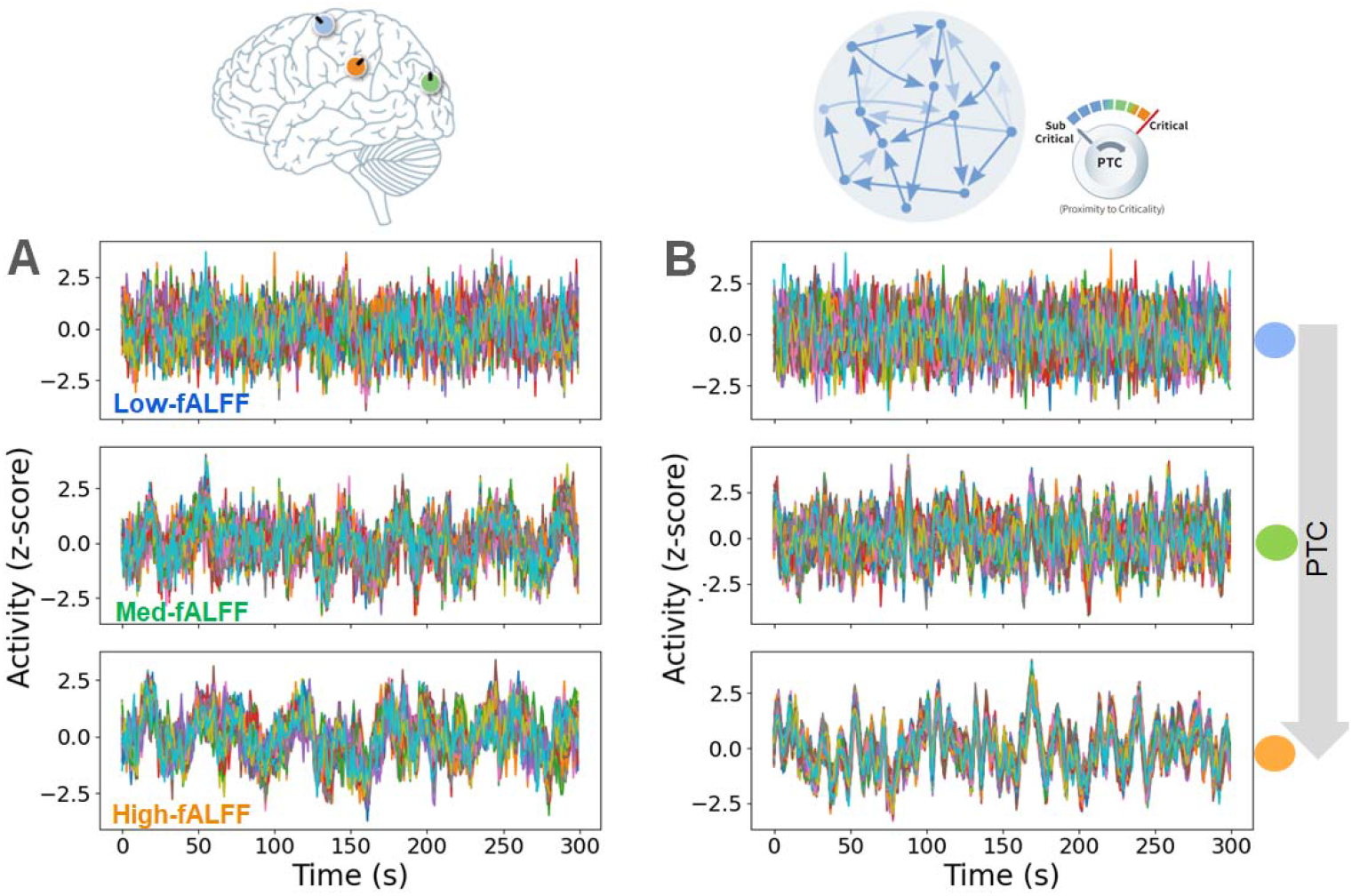
Distinct spontaneous oscillations dynamics in the brain and in a recurrent network simulation at different proximity to criticality. (A) Single-subject fMRI BOLD activity from voxels within representative cortical areas illustrating different fALFF levels: low (blue) in premotor FEF, intermediate (green) in higher-order visual MST, and high (orange) in the default mode network’s IPL. (B) Population activity from the random recurrent network model simulation in three different instances with increasing proximity to criticality. Note the diverse nature of cortical dynamics reflected in different areas and the ability of the model simulation to qualitatively match such dynamics through a small change (∼ 0.9 to 0.99) of single parameter – proximity to criticality.

Two aspects can be seen even by simple visual inspection. First, different cortical areas display visibly different fluctuation dynamics. In this example a cortical area belonging to the Default Mode Network (IPL region of the DMN; bottom of Figure 2A) showed increase in low frequency power compared to a premotor FEF cortex area (at the top of the panel). Second, changing the single parameter G of the artificial network was sufficient to statistically match this diversity in fluctuation dynamics. Interestingly, the change in G needed to produce this diversity was small, from ∼0.9 to 0.99.

To what extent did this diversity constitute a general cortical phenomenon? In Figure 3 we analyzed changes in fluctuation dynamics across the cerebral cortex using the fALFF metric. Figure 3A color codes the multi-subject mean fALFF, mapping the results for all 180 ROI s of the left hemisphere (LH). Our findings indicated remarkable dynamic variability, with roughly two-fold change across the cortex (∼0.3 to ∼0.6). I t should be noted that the very low mean fALFF values at the medial cortical margins (green colors) are likely artifactual, since these regions are known to suffer from BOLD susceptibility artifacts leading to low BOLD temporal SNR (tSNR) (Huck et al., 2023; Jamil et al., 2021; Murphy et al., 2007). However, filtering out ROI s with low tSNR (< 80) did not show any marked difference in the maps and other analyses (Figure S1, and Methods).

**Figure 3.**
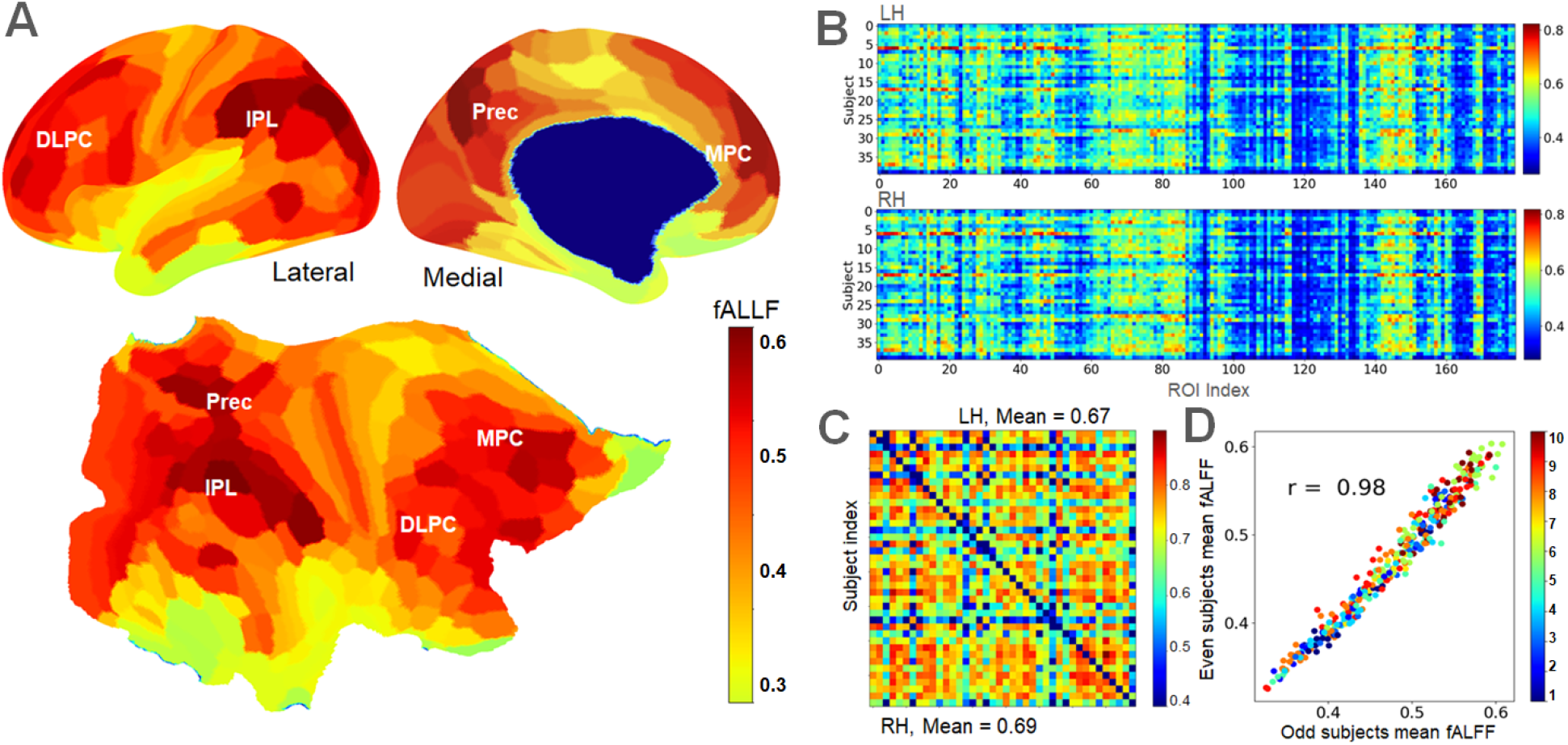
Spatial organization and reliability of cortical fALFF. **(A)** Multi-subject mean fALFF mapped onto the left-hemisphere cortex across 180 ROIs, revealing pronounced regional variability with approximately two-fold differences across cortical areas. **(B)** Distribution of regional fALFF values across individual subjects, demonstrating high consistency of fluctuation profiles across participants. **(C)** Pairwise correlations between per-subject fALFF distributions across all hemispheric ROIs, showing robust rank-order similarity across individuals (mean *r* ≈ 0.67–0.69 for LH and RH). **(D)** Split-half reliability analysis comparing population-averaged fALFF values computed from odd-versus even-indexed subjects across all ROIs, indicating high stability of the group-level fALFF profile.

In agreement with previous studies (Golesorkhi et al., 2021; Honey et al., 2012), high order regions, such as those related to the DMN, displayed the highest fALFF (i.e. particularly slow resting state fluctuations dynamics).

The sample size in this data set (N = 40) allowed us to examine the consistency of the regional fALFF scores across individuals. Panels 3B-D depict the results of this analysis. Ordering cortical ROIs according to their fALFF values revealed a striking cross-subject consistency, indicating that the rank-order of regions along this axis is highly preserved across individuals (Figure 3B).

Having noted this regional fALFF rank-order similarity across subjects, we compared all per-subject distribution pairings of the 180 hemispheric ROIs subject in Figure 3C indicating a mean correlation of r = [0.67, 0.69] in LH and RH, respectively. To gain insight into the stability of the population mean, the average ROI fALFF was computed for the odd vs. even indexed subjects across all ROIs. The result scatter is presented in Figure 3D, showing high correlation value across these two population averages (see related region-hierarchy color-coding and further details in Methods). Removal of low tSNR regions from the above analysis in Figure 3 had little influence on the statistical significance of the results, as shown in Figure S1. Comparing inter-subject variability in fALFF across cortical networks revealed lower variability within the DMN relative to other networks, though the variance explained remained highly significant across all regions (see Figure S2).

Next, we compared the empirical cortical fALFF distribution with that obtained from isolated recurrent-network simulations (Figure 4). As the simulated networks approached criticality, they naturally generated a range of fluctuation dynamics that overlapped with the empirical cortical fALFF values. Interestingly, when cortical ROIs were sorted according to their fALFF values, the resulting distribution was remarkably linear (*R*^2^ . 0.99; Figure 4A). Such near-linearity is expected when the underlying values are approximately uniformly distributed across their range (Methods).

**Figure 4.**
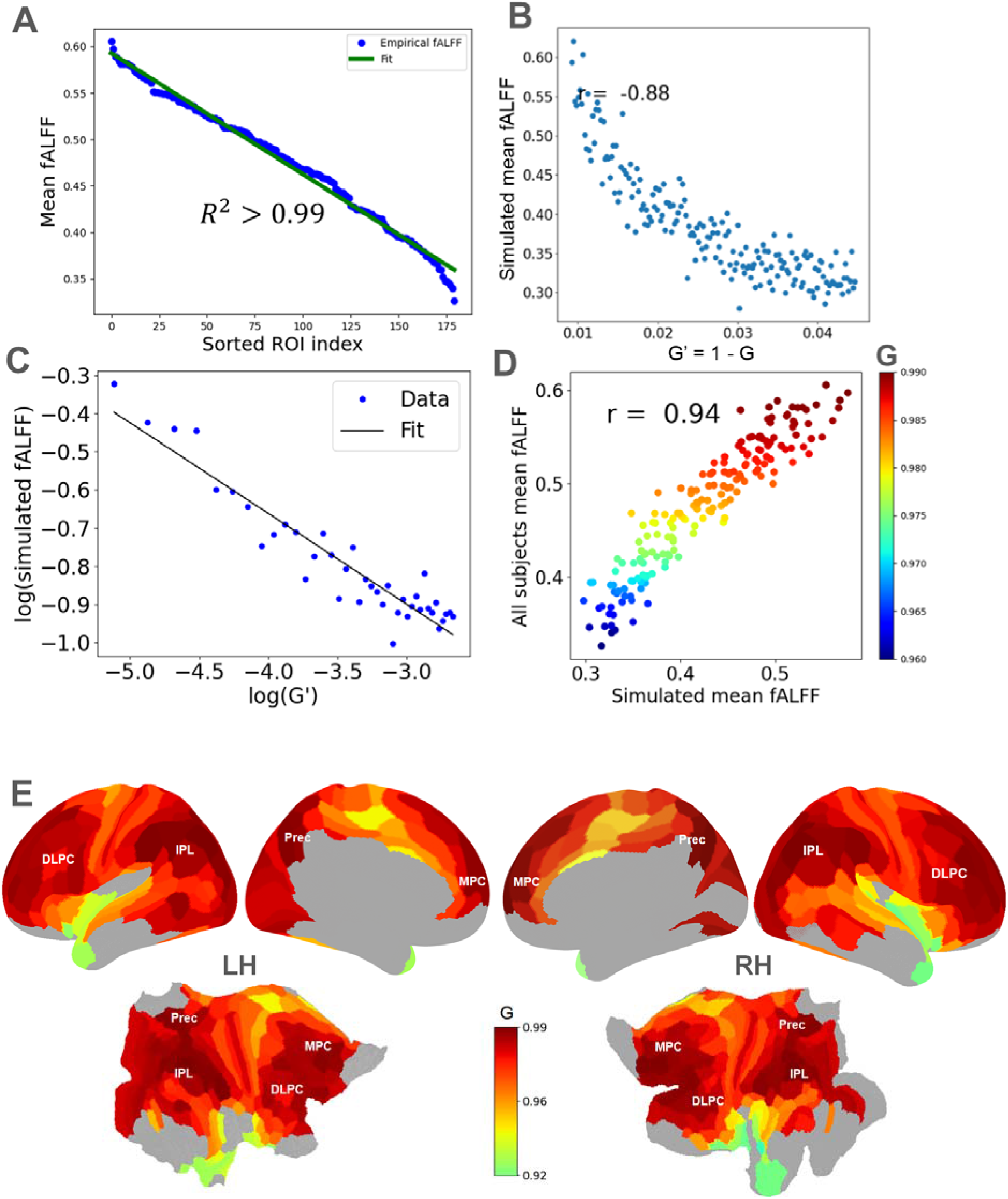
Empirical cortical fALFF suggests scale-free distribution in correspondence with local recurrent network simulations. **(A)** Empirical distribution of fALFF across cortical ROIs, showing an approximately monotonic linear slope, consistent with a near-uniform spread of regional fluctuation amplitudes. **(B)** Simulated fALFF obtained from random recurrent networks as neural gain, (note), is increased from sub-critical to near-critical regimes, reveals a pronounced non-linear dependence of fALFF on proximity to criticality. **(C)** Relationship between simulated fALFF and deviation from the critical point, demonstrating a power-law scaling. **(D)** Network simulations closely matched empirical fALFF across left-hemisphere ROIs over a log-spaced grid of neural gain values. Points indicate mean simulated fALFF across five realizations and are color-coded by *G*, supporting a power-law organization of cortical fALFF. **(E)** Map of distance to criticality across the LH and RH cortical sheet (tSNR-corrected), based on the estimated G.

In contrast, simulations in which neural gain was gradually increased from subcritical to near-critical values produced a strongly nonlinear relationship between fALFF and distance to criticality (Figures 4B and S3; for convenience we define G’ = (1-G)).

Specifically, the simulated fALFF followed an approximate power-law relationship with the distance from the critical point, consistent with the analytical prediction (Methods):

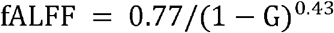

This result establishes a monotonic mapping between fALFF and proximity to criticality in the model. However, given the simplified nature of the recurrent-network model, we did not interpret this analytical relationship as providing a precise quantitative estimate of G for each cortical region. Instead, we used it to infer the overall distribution of proximity-to-criticality values across cortex. Specifically, the approximately uniform distribution of empirical fALFF values implies a corresponding logarithmic spacing of G values under the model’s power-law relationship. We therefore assigned logarithmically spaced G values to the rank-ordered cortical ROIs (Methods), yielding a robust estimate of the spatial distribution of proximity to criticality while avoiding over-interpretation of the exact model parameters.

Using this approach, the empirical and simulated fALFF values showed excellent agreement across all left-hemisphere ROIs (Figure 4D), supporting the hypothesis that cortical variability in fALFF reflects systematic variation in proximity to criticality. Importantly, the resulting cortical map should be interpreted as a model-based estimate of the relative distribution of distance to criticality across regions, rather than as a direct measurement of the underlying control parameter. The correspondence between empirical and simulated fALFF further improved when averaging across multiple network realizations (Figure S4). The spatial correlation increased from r = 0.81 for a single realization to r = 0.94 when averaging across five realizations, indicating increased stability of the estimated cortical profile.

Together, the whole-brain fALFF analysis and the corresponding recurrent-network simulations enable the construction of a cortical map of relative proximity to criticality (Figure 4E), grounded in the empirical spatial organization of resting-state dynamics and constrained by the model-derived relationship between fALFF and distance to criticality.

The fALFF measure is a single variable estimator of the fluctuation dynamics which does not capture the full profile of the power spectral distribution.

To obtain a more detailed characterization of the dynamics, we examined the full PSD across cortical regions and characterized its broadband aperiodic component in Figure 5. The aperiodic exponent, defined from the dependence of spectral power on frequency in log–log coordinates, quantifies the scaling of the spectrum; larger values indicate a stronger relative contribution of slow fluctuations. Panel 5A depicts the entire set of variance-explained BOLD PSD profiles (plotted on a log-log scale) obtained from all cortical regions, arranged from the regions showing the lowest to highest fALFF-based DTC estimation (color coded and in units of the control parameter G). To quantify spectral shape independently of overall power, singular-value decomposition (SVD) was applied to group-mean log-PSDs derived from demeaned time courses, after removing each region’s mean log power. A single component, PC1, accounted for 97.0% of cross-regional shape variance, whereas PC2 accounted for only 1.4%, with higher-order components explaining negligible additional variance (Figure 5B). Having identified a dominant spectral axis, we next asked whether this single mode could account for the apparent diversity. We tested this by centering the spectra relative to the shared axis and applying a gain-dependent rescaling (Methods). Figure 5C depicts the results of this shape collapse analysis aimed to rescale the data so that the distributions from different ROIs collapse onto a single universal curve, a hallmark of scale invariance (Ponce-Alvarez et al., 2023).

**Figure 5.**
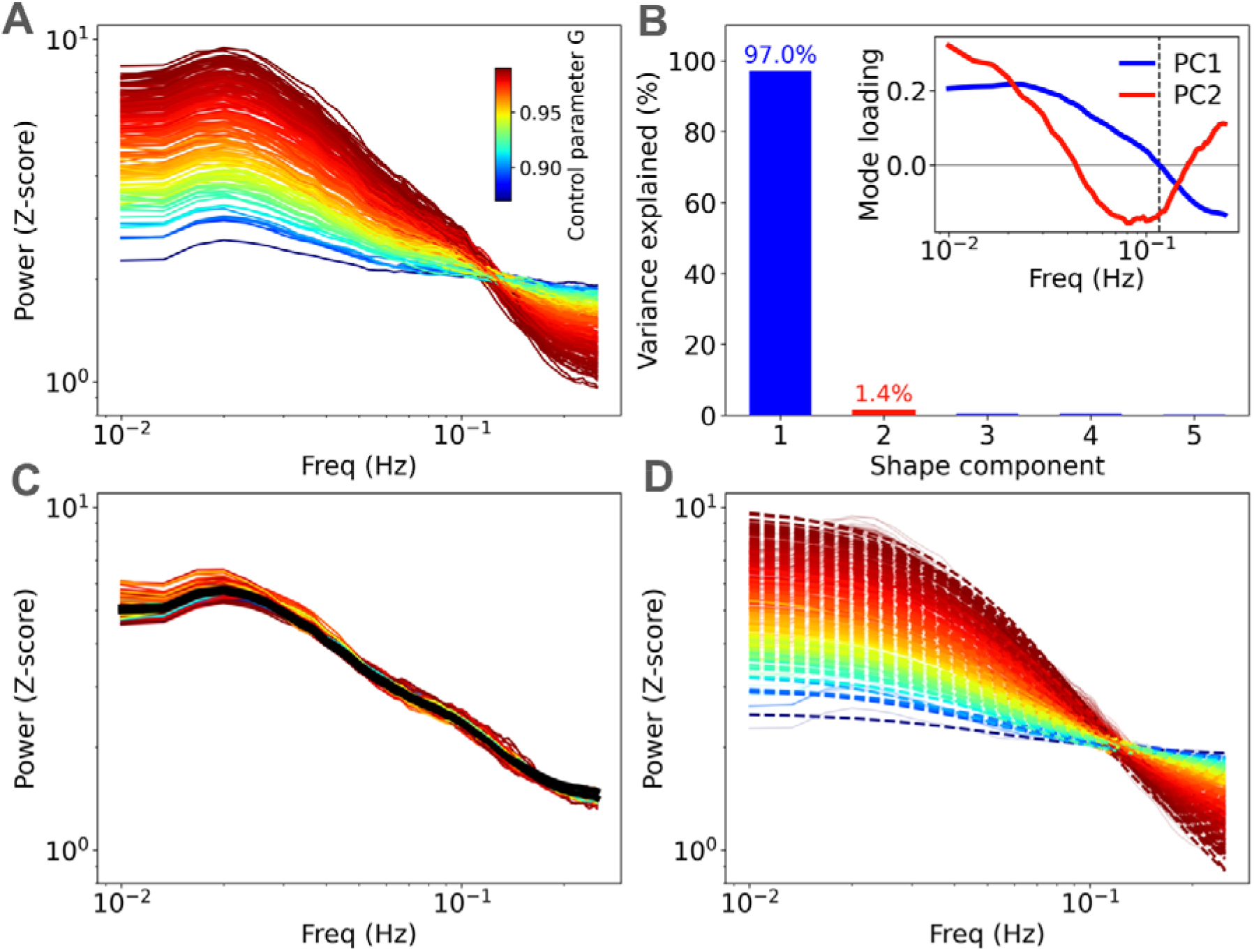
Power spectral density structure and shape collapse analysis. (A) PSD profiles of BOLD signals from all 180 cortical ROIs on log–log axes, color-coded by increasing fALFF-based DTC estimate. Note the apparent common vertex is a consequence of the variance normalization and is not interpreted as a characteristic frequency of cortical dynamics. (B) SVD of the regional log-PSDs after removing each region’s overall power, yields a single dominant component explaining 97.0% of the cross-regional shape variance (PC2 = 1.4%; components 3–5 negligible). Inset: the two leading shape modes — PC1 (blue) is a monotonic tilt whose zero-crossing (dashed, 0.117 Hz) corresponds to the crossing seen in (A), while PC2 (red) is a minor curvature mode. (C) Spectra were centered on the common PC1 crossing point and rescaled by a gain-dependent factor (Methods), demonstrating shape collapse to a single common curve (black, mean). (D) Analytical two-mode Lorentzian approximation of the cortical regional PSDs (dashed) overlaying the empirical profiles (background), capturing the overall empirical spectral shapes.

Given the relatively noisy single realization simulation-derived spectra, we first turned to an analytical two-mode Lorentzian approximation of random recurrent network PSDs (Chaudhuri et al., 2018; Yellin et al., 2025; see Methods and SI Appendix for details) to test whether the framework. This approximation captured the overall regional PSD shapes well (log-log .2 empirical cortical PSDs and their shape collapse could be reproduced within a reduced spectral median 0.99; Figure 5D), supporting similar shape collapse (98% of cross-regional shape variance; 86% reduction in cross-region spread; Figure S5A).

To establish the level of spectral similarity expected in the simulated network model, we first examined pairwise PSD similarity across cortical regions, which showed highly coherent spectral shapes across cortex (Figure S5B). In contrast, the spectral profiles obtained from single simulation realizations were substantially noisier (reflecting finite-size noise in the 400-unit recurrent networks), reaching comparable coherence levels to the empirical profiles only near the critical regime (G > 0.96; Figure S5C). Thus, demonstrating that increasing gain toward criticality reduces stochastic variability in the simulated dynamics and brings the model spectra closer to the empirical profiles. Due to the noisier profile in the subcritical range, PSDs were averaged across multiple simulation realizations prior to further analysis. Consistent with the analytical approximation, isolated recurrent-network simulations across DTCs closely matched and significantly correlated with the empirical PSDs (Figure S5D).

The other important aspect of the spontaneous fluctuations is their local correlation. In Figure 6 we examined rFCD (Methods), a metric of the regional pairwise correlation distribution, focusing on the location of the distribution peak. We hypothesize that the increased peak of the rFCD reflects converging DTC. Similarly to the fALFF distribution, the maps in Figure 6A show that the multi-subject mean rFCD peak also demonstrates remarkable diversity across cortical regions. Maps depict the distribution of rFCD peak values across 180 ROIs of the LH cortex. Low tSNR values of the rFCD, e.g. below 0.2, likely indicate high noise regions associated with BOLD susceptibility artifacts (see tSNR corrected maps and analysis in Figure S6). As in the fALFF analysis the dataset size was well suited to directly measure of the reliability of the rFCD peak across participants. Panels 6B-D analyze the comparison across individuals and ROIs, revealing a highly preserved rank-order sorting of ROIs as a function of rFCD peak, resembling the same organization shown by the fALFF measure. As shown in Figure 6B, ordering cortical regions by rFCD values revealed a highly consistent rank-order across individuals, closely mirroring the pattern observed for fALFF. Correlation of rank-order pairings across the 180 hemispheric ROIs (Figure 6C) yielded mean values of r = [0.58, 0.57] for LH and RH, respectively—slightly lower than the corresponding fALFF scores. Furthermore, analysis of odd-versus even-indexed subject groups across all ROIs (Figure 6D, see color-code of data points in Methods) revealed a particularly strong correlation between the two population averages, underscoring the stability of the mean rFCD distribution (again resembling the fALFF results).

**Figure 6.**
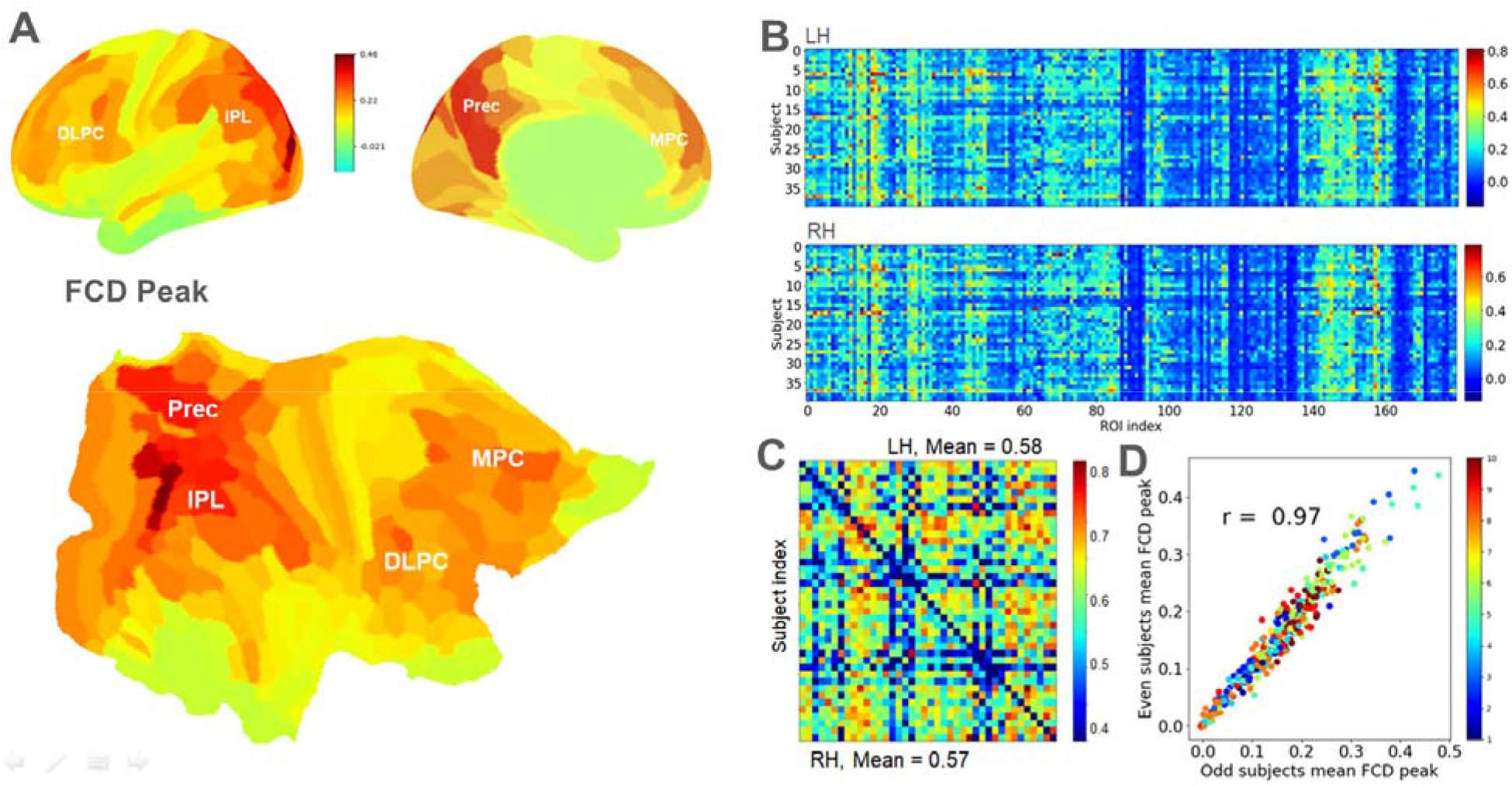
Spatial organization and reliability of regional functional connectivity density. **(A)** Multi-subject mean rFCD peak mapped onto the left-hemisphere cortex across 180 ROIs, revealing substantial regional variability in local correlation structure. Lower rFCD peak values (e.g., < 0.2) are predominantly observed in regions with reduced BOLD temporal SNR and are likely influenced by susceptibility-related noise. **(B)** Distribution of rFCD peak values across individual subjects, indicating consistent regional profiles of local functional connectivity across participants. **(C)** Pairwise correlations between per-subject rFCD peak distributions across all hemispheric ROIs, indicating a highly preserved rank-order organization of regions across individuals. **(D)** Split-half reliability analysis comparing population-averaged rFCD peak values computed from odd-versus even-indexed subjects across all ROIs, showing strong stability of the group-level rFCD organization and a regional ordering closely resembling that observed for fALFF.

To what extent does rFCD echo similar patterns previously identified with fALFF, and can network models shed more light on the underlying mechanisms of this relationship? We address these questions in Figure 7. As in the spontaneous fluctuations’ power spectra, the profile of regional correlations showed a substantial diversity across different cortical regions. Figure 7A depicts the distribution of these profiles across the entire set of cortical ROIs. Here, we transform rFCD histograms into kernel density estimate (KDE) plots (Methods). KDE provides a smoothed, continuous representation of the distributions, allowing more robust cross-regional comparison while reducing sensitivity to variability in voxel sampling across ROIs.

**Figure 7.**
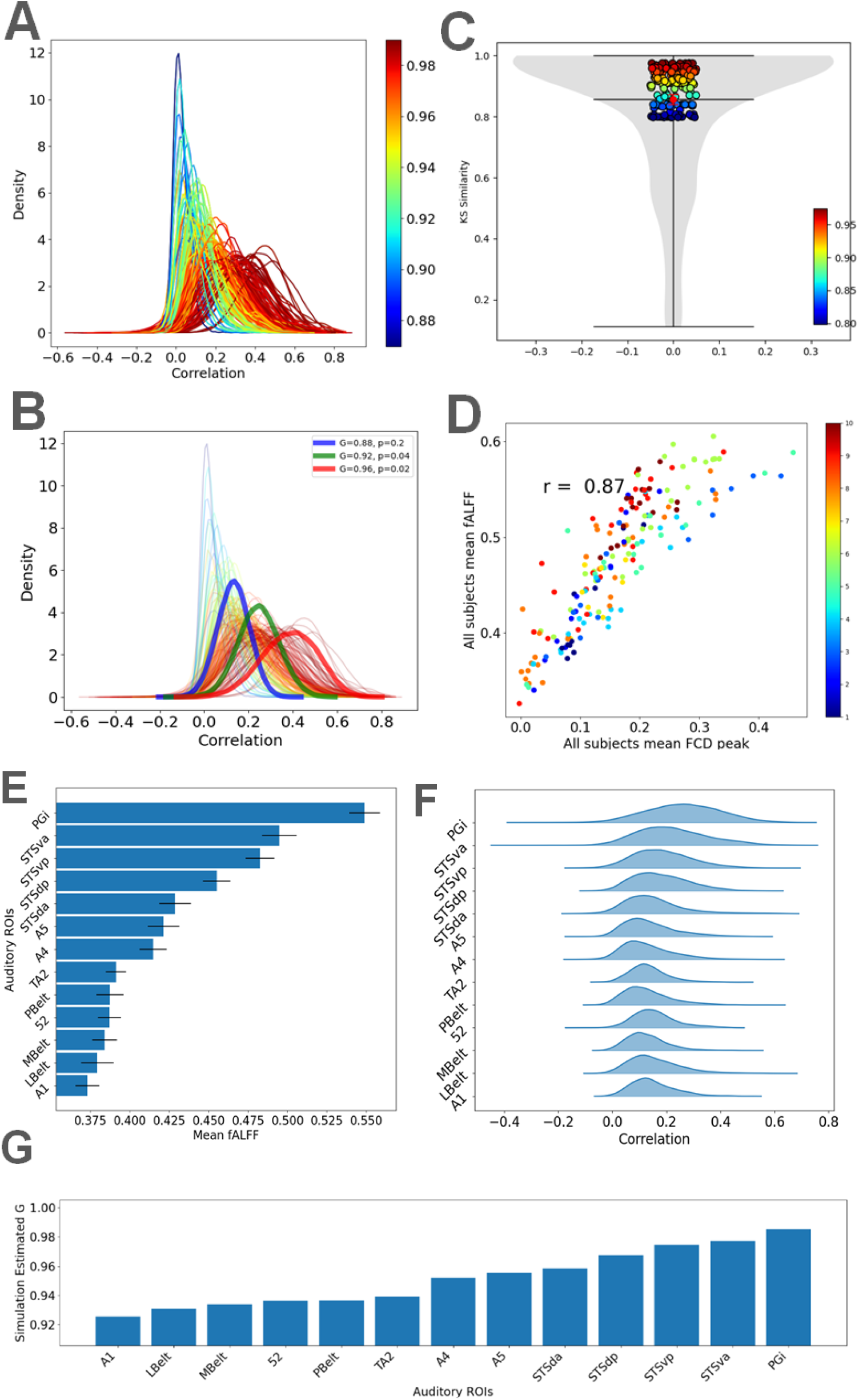
Regional functional connectivity density (rFCD) diversity, model correspondence, and hierarchical organization. **(A)** Kernel density estimate (KDE) representations of rFCD distributions across all cortical ROIs, revealing substantial regional diversity in correlation profiles. **(B)** Representative examples of simulated rFCD distributions overlaid on the empirical BOLD-fMRI rFCD profile, demonstrating close correspondence between data and isolated-network simulations at different combinations of neural gain (*G*) and connectivity sparseness (*p*). **(C)** Goodness-of-fit between empirical and simulated rFCD distributions across all cortical regions, quantified using Kolmogorov–Smirnov (KS) similarity (mean ± SEM). As with fluctuation power spectra, optimal correspondence is achieved by tuning proximity to criticality, with best overall fits obtained by adjusting network sparseness. **(D)** Across-subject spatial correlation between regional rFCD peak values and fALFF, indicating that regions with stronger local connectivity also exhibit slower intrinsic fluctuations, consistent with their previously reported temporal coupling. **(E–G)** Hierarchical organization of DTC metrics along the auditory cortical hierarchy. **(E)** Mean fALFF increases progressively from early to higher-order auditory regions. **(F)** rFCD distributions exhibit a gradual rightward shift toward higher mean correlation values together with increased distribution width, consistent with slower intrinsic timescales. **(G)** Corresponding increases in estimated neural gain (*G*) across the hierarchy. Auditory regions, sorted along the hierarchy, included: A1 → LBelt/MBelt/52 → PBelt → A4 → A5 → STGa/STSda/STSdp → TPOJ1/STV → PGi.

Figure 7B portrays the typical fit matches of few example simulation realizations on top of the empirical rFCD distribution. We observed that the network sparseness parameter p exerted a substantial influence on the simulated rFCD distributions. Decreasing p (i.e., increasing sparsity) led to a progressive widening of the correlation distribution curves. Although the initial simulations used p = 0.2, optimal fitting of near-critical ROIs necessitated a markedly sparser configuration (p <≈ 0.04). As can be seen, the BOLD-fMRI correlation profile and the simulated network ones were well matched, at G = 0.88 & p = 0.2, marked in blue; G = 0.92 & p = 0.04 in green; and G = 0.96 & p=0.02 in red).

To quantify the correspondence between the simulations and the empirical BOLD-fMRI correlation profile, we computed a similarity measure between the two distributions. Figure 7C shows the goodness of fit values (using KS similarity mean fit and SEM scores; Methods) of the simulated rFCD relative to all cortical regions. Similarly to the case of the power spectra distributions, the fit to the entire set of cortical regions was obtained by manipulating a single parameter: the closeness to criticality. Kolmogorov-Smirnoff (KS) based similarity ranking indicated significant fit for the above. However, both the simulations and the corresponding bar-plot comparisons indicated that optimal fitting required adjusting the connectivity sparseness to p=0.04.

Figure 7D demonstrates the individual subject rFCD peak to fALFF correlation scatter, a spatial correlation of these two measures echoing their previously found temporal relation, notably within distributed association networks (Di et al., 2013; Tomasi et al., 2016; Tomasi & Volkow, 2019). The observed rFCD distribution may reflect multiplicative signal propagation in a sparse network operating near criticality, naturally yielding a skewed, approximately lognormal profile of pairwise correlations. A formal evaluation of this hypothesis, however, lies beyond the scope of the present study and is reserved for future work.

Interestingly, when we examined the DTC within known cortical systems, it became apparent that a sequential hierarchy can be detected. An example of this phenomenon is depicted In Panels 7E-G, where we focus on the Auditory hierarchy and show how all metrics indicate a gradual increase from early to high-order areas, suggesting a monotonic increase in proximity to criticality along this hierarchy. Figure 7E presents a consistently growing mean fALFF. Figure 7F shows how the center of rFCD gradually right-shifts (to higher level of mean correlation) and how the KDE distribution widens (implying slower timescale). Our analysis confirms previous reports of cortical time-scale hierarchies (Lerner et al., 2014; Raut et al., 2020), suggesting DTC as the underlying mechanistic principle. Finally, Figure 7G presents the increase in estimated G. The analysis included the following Auditory hierarchy regions: A1 ->⍰LBelt, MBelt and 52 ->⍰PBelt -> A4⍰ -> A5⍰ -> STGa, STSda, STSdp -> TPOJ1, STV -> PGi.

## Discussion

Our study explores the hypothesis that a substantial part of cortical functional diversity may be explained by a single underlying dynamical mechanism whose tuning gives rise to a wide range of behaviors. We identify distance to criticality, a fundamental dynamical property of recurrent cortical networks, as a unifying mechanism underlying the functional diversity observed across human cortical areas. Multiple types of critical transitions have been proposed in neural systems, including transitions to chaotic and pattern-forming regimes (O’Byrne & Jerbi, 2022b). Here, we focus on instability criticality, characterized by a transition between stable and unstable activity, which we probe by modulating neural gain in a random recurrent network.

Using resting-state fMRI data from the Human Connectome Project (Glasser et al., 2013), comprising 40 participants scanned at high field (7T), we demonstrate that variations in the distance of local networks from the critical point accounts for two central features of cortical dynamics: the power spectra of ultra-slow spontaneous fluctuations observed across cortex at rest (Fox & Raichle, 2007; Nir et al., 2006, 2008), and the functional connectivity profile (Biswal et al., 1995; Fox et al., 2006; Harmelech & Malach, 2013), focusing on the regional cross-voxel correlation structure across cortex.

Our analysis of the HCP data confirms previous reports that both measures vary substantially across cortical regions (Tomasi & Volkow, 2011; Yang et al., 2007). The relative power of the ultra-slow fluctuations (fALFF) in our analysis ranged from ∼0.3 to ∼0.8 across the cortex, excluding low SNR regions. Similarly, our analysis of the peak in local correlation structure (rFCD) showed values that ranged from ∼0.1 to ∼0.6, again excluding the low SNR regions.

By fitting a recurrent network model to the data, we demonstrate that DTC is spatially varying across the resting human cortex, with regions occupying a cross-individual stable and highly conserved rank order along this axis. Importantly, this organization reflects marked differences in DTC across cortical regions, indicating that, during rest, some networks operate systematically closer to the transition regime while others remain further removed.

This orderly distribution suggests that proximity to the transition regime represents an intrinsic, system-level organizational principle, that coexists with previously demonstrated transient shifts in DTC. Such transient shifts have been observed across a range of behavioral and physiological conditions. For example, proximity to criticality has been shown to increase during free-recall tasks (Yellin et al., 2025), as well as during prolonged wakefulness (Meisel et al., 2015), where it correlates with behavioral performance such as reaction time (Harel et al., 2025). Conversely, pathological states such as epileptiform activity are associated with deviations toward supercritical dynamics (Arviv et al., 2016), highlighting the sensitivity of cortical dynamics to both functional demands and disease processes.

Critically, our results demonstrate that much of this rich diversity can be captured by manipulating a single parameter, namely the neural gain (as part of the criticality control parameter G), in a simple network of random recurrent connections with varying DTC. Realistically, a highly simplified model such as our small recurrent random network cannot be expected to encompass the multi-dimensional complexity of a vast biological neuronal network. However, as shown in Figure 4, manipulating neural gain alone was enough to fit all the fALFF data obtained in the diversity of cortical regions along the dimensions of power spectra and local connectivity. This outcome is remarkable if one considers that the dynamic input that was fed into the network simulation was drastically different from the output of the network: first, its frequency profile (PSD) was flat-consisting of random “white” noise (see Methods). Second the inputs to each artificial neuron were strictly independent. As we have argued elsewhere (Yellin et al., 2025), the general slowing down of the network dynamic output-despite its fast random inputs can be attributed to a fundamental property of recurrent networks, termed critical slowing down.

Regional cortical PSDs were organized predominantly along a single broadband spectrally-scaled dimension, which accounted for 97% of cross-regional shape variance and closely corresponded to the aperiodic exponent. Rescaling each region’s projection onto this dominant component produced a marked PC1-based shape collapse, indicating that regional differences are largely described by movement along a common one-parameter spectral family. Because the aperiodic exponent and fALFF capture closely related aspects of the same redistribution of spectral power, their associations with the inferred DTC should not be regarded as independent evidence. The small residual variance may contain additional structure, including low-frequency curvature, although whether this reflects a distinct biological timescale remains unresolved.

More broadly, the present simulations showed that recurrent slowing down reproduced not only the overall profile of relative power changes across ultra-slow fluctuation frequencies (< 0.1 Hz) observed in the human BOLD response (Figure 5), but also the profile of within-region voxel correlation levels reflected in the FCD (Figures 6 and 7). Together, these findings suggest that variation in distance to criticality can account for multiple empirical signatures of cortical dynamics within a common framework, spanning both the spectral organization of regional fluctuations and the local correlation structure of voxel-level activity.

While it could be argued that other, more complex models, may also be able to successfully match this rich data set, the point of this study was to demonstrate that such diverse repertoire of dynamics and connectivity can be matched by a simple, recurrent network, through the modulation of a single parameter - proximity to the edge of dynamic instability. This strongly supports the notion that the dense network of local connections, one of the central features of cortical micro-architecture (Amir et al., 1993; Malach et al., 1993; V. Mountcastle, 1997; V. B. Mountcastle, 1957) as well as cortical neuronal models (Wieland et al., 2015), plays a pivotal role in generating the observed neuronal dynamics in the cerebral cortex. Furthermore, it supports the notion that an instrumental gain control parameter underlying the rich dynamics is controlling the proximity to criticality in these networks.

What could be the function of controlling the proximity to criticality in the cortex? An attractive hypothesis argues that transient short crossing of the critical point, and the ensuing phase transition associated with such crossing, underlies cortical decisions, across diverse modalities. For example, in the sensory domain, the non-linear “ignitions” associated with such transition underlies perceptual, awareness related decisions (Deco & Jirsa, 2012; Fisch et al., 2009; Moutard et al., 2015; Shriki et al., 2013). We have previously proposed that by transiently manipulating DTC, the cortex can control the ease by which networks can reach their decision thresholds (Norman et al., 2017; Yellin et al., 2025). It is interesting in this respect to note that during rest, the network that was estimated to be closest to the transition threshold was the DMN. This is nicely compatible with the observations that DMN activity can be easily modulated during no-stimulus condition (Harmelech et al., 2013; Honey et al., 2012; Murray et al., 2014; Tagliazucchi et al., 2012; Yellin et al., 2015). This also fits well with findings indicating that the resting state is associated with spontaneous recollection and planning events (Kucyi et al., 2016), all suggesting that the DMN is more prone to crossing the criticality threshold during rest compared to other networks, as was shown through our network simulation, in the present study. Together, these findings are consistent with the view that higher-order networks operate closer to criticality, supporting internally generated and temporally extended cognitive processes (Algom & Shriki, 2026).

A natural interpretation of these converging findings is that the preferential positioning of DMN regions near the transition threshold in our cortical map provides a mechanistic substrate for their role in long-timescale processing (Honey et al., 2012; Lerner et al., 2014). Operating at minimal distance to criticality, DMN circuits would be expected to exhibit enhanced temporal integration, slow intrinsic fluctuations, and heightened sensitivity to structured inputs - precisely the dynamical features required for tracking extended narrative structure and accumulating information over long periods. In this framework, the DMN’s proximity to the critical point may not merely reflect a static architectural trait, but rather a dynamical configuration that supports amplification and integration of temporally extended content.

A potential concern is that the observed spatial distribution of fALFF and rFCD could in part reflect vascular (e.g. venous) rather than neuronal contributions. However, Huck et al., 2023 demonstrated that while ALFF is moderately affected by venous bias, fALFF shows an almost negligible effect (≈0.03% change across cortex) due to its ratio normalization, a finding consistent with (Vigneau-Roy et al., 2014), who likewise reported minimal vascular influence on fALFF maps. These results collectively indicate that the present findings are unlikely to be driven by venous artifacts.

To mitigate potential noise-related confounds, Harris et al., 2024 proposed a noise-robust DTC metric known as rescaling of autodensity (RAD), designed to reliably track DTC even under fluctuating noise amplitudes (Liu et al., 2025). When applied to mouse electrophysiology data, RAD revealed, consistent with our own findings, that higher-order visual regions are closer to criticality than lower-order areas, suggesting a hierarchical organization in the brain’s progression toward critical states.

Whereas fALFF values across cortical ROIs followed a scale-free spatial distribution, the rFCD distribution followed a markedly different, broad and skewed form. One possible mechanistic interpretation of this distinct rFCD pattern is the multiplicative nature of signal propagation in sparse recurrent networks operating near criticality. Pairwise functional correlations can be viewed as the aggregate contribution of many multi-step interaction pathways, each defined by a product of synaptic weights. Because products of random variables tend toward lognormality, such multiplicative path structure naturally predisposes the distribution of effective coupling strengths to be highly skewed (Buzsáki & Mizuseki, 2014). Operating near criticality further amplifies this effect: marginally stable dynamics allow longer pathways to remain influential, compounding small stochastic deviations across many steps and yielding a heavy-tailed distribution in which most neuron pairs are weakly coupled while a small minority exhibit disproportionately strong correlations. Analogously, in Kolmogorov’s multiplicative-cascade account of turbulence intermittency (Yamazaki, 1990), repeated scale-to-scale amplification produces lognormal-like variability, suggesting that near-critical cortex-wide slow-mode fluctuations may similarly cascade across network pathways to shape the skewed rFCD distribution. Although this framework provides a plausible explanation for the observed rFCD profile, a formal analytical and simulation-based characterization of this multiplicative, near-critical mechanism will be pursued in future work.

In summary, we demonstrate here that the rich repertoire of resting-state fluctuation dynamics and correlations found across the entire set of cortical regions can be accounted for by a simple recurrent network model, through controlling a single parameter - proximity to criticality. The findings highlight the central role of local recurrent circuitry and critical dynamics in shaping cortical function. More broadly, they extend a growing body of work suggesting that proximity to criticality provides a unifying framework linking neural dynamics across scales, conditions, and functional domains (Fekete et al., 2018; Maturana et al., 2020).

## Methods

### The human connectome project fMRI data and its analysis

Functional MRI data were obtained from the Human Connectome Project (HCP) Young Adult dataset and were processed using the HCP minimal preprocessing pipelines (Glasser et al., 2013). All HCP data were collected with written informed consent from participants under protocols approved by the relevant Institutional Review Boards, and in accordance with the principles of the Declaration of Helsinki. This study analyzed only de-identified publicly available data. Briefly, functional images underwent gradient-distortion correction, motion correction, and susceptibility-induced distortion correction using reversed phase-encoding acquisitions. Functional images were registered to each subject’s structural T1-weighted image using boundary-based registration and resampled to standard grayordinate space. Cortical time series were projected onto the cortical surface using ribbon-constrained volume-to-surface mapping, while subcortical signals were retained in volumetric form and combined with surface data into CIFTI dense time-series (.dtseries.nii) files. Cross-subject surface alignment was performed using the MSMAll multimodal surface matching algorithm (Glasser et al., 2016). The time series were high-pass filtered (cutoff = 2000 s) and denoised using ICA-FIX. The resulting cleaned grayordinate time series (*_Atlas_MSMAll_hp2000_clean.dtseries.nii) were used for subsequent analyses.

### fMRI data Parcellation

To reduce the dimensionality of the data and facilitate connectomic analyses, we parcellated cortical rs-fMRI time series using the multimodal cortical atlas of (Glasser et al., 2016), which subdivides the cerebral cortex into 180 structurally and functionally defined regions of interest (ROIs) per hemisphere. The Glasser parcellation combines multimodal MRI features to define boundaries of cortical areas and has been widely adopted in HCP-based connectivity studies.

### Hemodynamic response function (HRF) convolution

To enable comparison with empirical resting-state fMRI signals, the simulated neuronal firing rate time series generated by the dynamics model were convolved with a canonical hemodynamic response function (HRF). This step accounts for the temporal delay and smoothing inherent to neurovascular coupling, thereby producing model-based BOLD signals that could be directly compared with the spontaneous activity observed in the empirical data.

Recent work further demonstrates that resting-state fMRI signals contain spectral signatures of local HRF timing differences, highlighting the importance of explicitly modeling the hemodynamic transformation when linking neuronal dynamics to BOLD measurements (Bailes et al., 2023).

### Random recurrent network as a rate model

We simulated a random recurrent network model consisting of N rate-based units representing single neurons or groups of neurons. We closely follow the model proposed in ( Yellin et al., 2025 and Chaudhuri et al., 2018) to study DTC metrics in HCP fMRI data. The firing rate dynamics of the j-th neuron’s output over time in this model is given by:

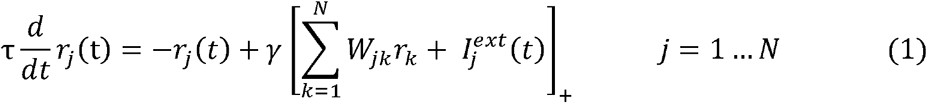

where *r_j_* is the firing rate of the *j*-th unit activity level, and *τ* is a characteristic integration time. The [ ] _+_ brackets denote a rectified linear (ReLU) activation function. The *γ* variable denotes the slope of the firing rate – current curve (f-I curve). The weights *W_jk_* describe the strength of the connection from the presynaptic unit k to the postsynaptic unit *j*, and 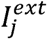 denotes the external input to the j-th unit (drawn here randomly from a “standard normal” distribution).

### Recurrent rate model ofROI-level fMRI voxels

The present model is formulated at the level of firing rates, abstracting away individual spikes and synaptic conductance dynamics. This approximation is well justified when modeling large, recurrent cortical populations operating in an asynchronous regime, where collective dynamics are dominated by slowly varying population activity rather than precise spike timing. In such regimes, spiking networks of conductance-based neurons can be approximated by low-dimensional rate models that capture the dominant temporal modes of the network dynamics (Shriki et al., 2003). Moreover, spiking-network models exhibit the same critical transition dynamics, showing maximal variability at the edge of instability, demonstrating that such critical regimes are preserved beyond rate-based abstractions (Karimipanah et al., 2017). Importantly, the fMRI BOLD voxel-level signal reflects temporally smoothed, population-averaged neural activity at spatial scales encompassing tens of thousands to millions of neurons and temporal scales of seconds. Rate-based descriptions therefore provide a natural and parsimonious link between underlying recurrent network dynamics and macroscopic observables measured with fMRI, while retaining the essential dynamical features relevant for critical slowing down and long-range temporal correlations.

### Criticality in the random recurrent network

We consider a network architecture comprising a non-hierarchical cluster of recurrently connected neurons characterized by a semi-linear f-I curve with gain γ. Following (Chaudhuri et al., 2018), we choose the connections in this network (the entries of matrix W) to be sparse and random. More specifically, each entry is nonzero with probability p, and the nonzero entries are drawn from a normal distribution, with positive mean μ_conn_ and variance σ_conn_, normalized by the size of the network, N.

The mean and STD of .ext, the noisy input, was ∼20 (±11.5) *p*A

The sampling fraction, α, for computing the signal, was set to1.0.

To provide additional insight in the present context, we note that near a steady state the spontaneous fluctuation dynamics can be approximated by a linearized algebraic system. We briefly summarize the key results (Yellin et al., 2025). The eigenvalue spectrum of the resulting random connectivity matrix consists of a bulk cluster of complex eigenvalues with negative real parts and a single isolated eigenvalue corresponding to the uniform eigenvector (i.e., the mean eigenvalue, λd =-1+G., which approaches zero as the system nears the bifurcation point at network activity). The matrix governing the linearized dynamics therefore has a dominant *G*=1, marking the transition from stable to unstable dynamics. This analysis relies on random matrix theory and thus assumes a large-scale network (Rajan & Abbott, 2006; Tao, 2013).

### Odd-vs-even analysis

The analysis in Figures 3 and 6 for fALFF and rFCD metrics separated the subject population into two halves by odd and even indexes. For each sub-population we averaged the metric value, for each ROI. The outcome scatter consists of plot where the x- axis shows mean value for the even group and the y-axis for the odd group. I.e. each ROI represents a single point in the scatter plot. Data points in the scatter were color-coded according to the following scheme. Given the 22 cortical systems (“cortical_ID”) defined by the anatomical grouping distributed with the HCP-MMP atlas (Glasser et al., 2016); we further consolidated this labeling into 10 hierarchical cortical classes spanning lower-order sensory regions to higher-order association cortex. This hierarchical grouping was motivated by known large-scale gradients of cortical organization, spanning unimodal sensory cortex to transmodal association networks. The mapping from the 22 cortical systems to the 10 hierarchical classes was as follows: (1) primary sensory and motor (1,10); (2) secondary sensory (2,11); (3) mid-order sensory and premotor (3,8); (4) high-order sensory (4,12,14); (5) early association (5,15); (6) attention (6,16,17); (7) complex motor planning (7); (8) default mode network (13,18,19); (9) frontoparietal (9,20); and (10) high-order cognitive (21,22). Analyses were performed at the parcel level, with statistical summaries computed across parcels belonging to the same hierarchical class.

### Scale-free distribution analysis and mathematical derivation

Assuming the underlying distribution is uniform, every value within the range is equally likely. When we sort such values from largest to smallest, we are essentially dividing the interval between the maximum and minimum into equal steps. Each ranked value will then be spaced at nearly constant intervals, producing a straight line when plotted against rank. In contrast, non-uniform distributions (such as Gaussian or exponential) would show curvature—values would cluster near one end and the rank plot would deviate from linearity. The near-linear falloff of the fALFF values therefore indicates that the regional amplitudes are spread almost evenly across their range, consistent with a uniform distribution.

The analysis in Figure 4 was based on the following steps:

a. sorting and indexing the empirical fALFF from large to small
b. collecting fALFF values from localrandom recurrent networks with linear gradual incresing G.
c. yielding: A = 0.06,α= 0.5 => leading to a prediction that:*f ALFF =A/(1-G)*^α^ fitting power-law parameters to estimate fALFF dependency on the control parameter G, f = linspace(0.3, 0.6, 180) => G_grid_ =1-(*A/f)^1/α^*
d. To generate the G-grid for the simulations of 180 LH cortical regions, we used:

### Spectral analysis for empirical–simulation PSD comparison

For each signal, the PSD was estimated using Welch’s method. Population differences in absolute spectral power were evaluated using the raw PSDs. To compare spectral shape independently of overall power, spectra were represented in log-log space and parameterized as: log(*P*(*f*)) =*β*0+*β*1log *f* + *β*2(log *f*) 2 + *ε*(*f*), where, *β*0 denotes broadband offset, *β*1 denotes spectral slope, and *β*2 denotes curvature. Group comparisons were then performed separately on the fitted offset, slope, and curvature parameters.

### Spectral dimensionality and PC1-based shape collapse

To quantify the dimensionality of regional spectral variation independently of the overall signal amplitude, we analyzed PSDs computed from temporally demeaned voxel time courses. Group-mean regional PSDs were log-transformed, and each region’s mean cross-frequency log power was removed to isolate spectral shape from broadband amplitude differences. The resulting region-by-frequency matrix was centered across regions and decomposed by singular-value decomposition (SVD). The first component accounted for 97.0% of cross-regional shape variance, whereas higher-order components contributed negligibly. Regional PC1 scores were then compared with the fitted aperiodic exponent, identifying PC1 as the dominant broadband spectral-scaling mode.

To visualize this one-dimensional organization in the variance-normalized spectra, each regional log-PSD was projected onto the fixed PC1 mode and its region-specific PC1 contribution was removed. The residual spectra were then restored around the common mean spectral profile and transformed back to linear power units, producing a PC1-based shape collapse. Similar result was obtained by scaling the spectra with a fitted function of the fALFF-derived DTC estimates. However, unlike this fit-based approach, the PC1 procedure required neither DTC estimates nor an additional scaling parameter; it therefore provides a direct visualization of the one-dimensional spectral organization identified by the SVD.

### Kernel Density Estimation of rFCD Distributions

Here, we transform rFCD histograms into kernel density estimate (KDE) plots. KDE is a method for normalizing distributions. In terms of data visualization, a histogram presents the frequency of data values falling within predetermined intervals (also known as “bins”). In contrast, a KDE plot illustrates the density estimate of the data values as a continuous curve. Based on a standard KDE implementation in Python’s SCiPy library, we implemented the following demonstration of the FCD-KDE spread along the Auditory hierarchy (according to the definition we used when plotting fALFF for this same hierarchy). Note ROIs are of different size (i.e. each having a different number of voxels), whereas the simulated network is of constant size (400 units). Moving to KDE naturally provides us normalized distributions to be compared.

## Declarations

## Ethics approval and consent to participate

This study involved secondary analysis of de-identified publicly available data from the Human Connectome Project (HCP). All HCP participants provided written informed consent, and data acquisition was approved by the Washington University Institutional Review Board in accordance with the Declaration of Helsinki. No additional ethics approval or participant consent was required for this study.

## Data availability

The datasets analyzed during the current study are publicly available from the Human Connectome Project (HCP; Young Adult S1200 release) to qualified investigators subject to the HCP data use terms. All data used in this study can be obtained from the HCP website.

## Code availability

The analysis and simulation code is available at: https://github.com/dovi-yellin/hcp_dtc

## Author contributions

D.Y., R.M. and O.S. conceived the study. E.S. preprocessed data. D.Y. designed and performed the data analysis, developed and executed software simulations, and prepared the figures. R.M. and O.S. supervised the project. All authors contributed to the writing, critically revised the manuscript, and approved the final revision.

