## Supplementary Information for "A cortical gradient of distance to criticality governs large-scale resting-state fMRI dynamics"

**Introduction**

Here, we provide additional analyses and figures supporting the main text results on cortical fALFF and rFCD, together with their corresponding random recurrent network simulations. The SI figures further test the reliability of the link between resting-state dynamics and distance to criticality (DTC), including its signal-to-noise sensitivity and inter-subject variability. Additional analyses characterize the behavior of large-scale recurrent network simulations across an extended range of neural gain values, revealing non-linear effects on slow-timescale dynamics and improved correspondence with empirical data when averaging across realizations. Further spectral analyses quantify the agreement between simulated and empirical power spectral densities and extend the shape-collapse results. Finally, subject-wise analyses of the rFCD–fALFF relationship further extend the reported link between local functional connectivity and spectral profile. These analyses provide additional support for the central claim that proximity to criticality governs both spectral and correlation structure in cortical dynamics.

**Fast and slow modes in random recurrent neural networks**

In both biological and artificial *random* recurrent networks, a useful technical view is that “fast” and “slow” activity reflects different spectral components (modes) of the network’s linearized dynamics around a working point (Chaudhuri et al., 2018; Rajan & Abbott, 2006; Tao, 2013; Yellin et al., 2025). A formal analysis of fast and slow activity in random recurrent networks is typically obtained by linearizing the dynamics around a fixed point of the network equation:

$\tau ẋ(t) = -x(t) + W \varphi(x(t)) + \eta(t)$	(1)

where, $x\left( t \right)\in\mathbb{R}^{N}$ is the vector of population firing rate, and τ is the intrinsic time constant governing passive decay $\left( -x \right)$. $W \in\mathbb{R}^{\left\{ N\times N \right\}}$ is the recurrent connectivity matrix, where $W_{ij}$ specifies the synaptic weight from unit $j$ to unit $i$. $\varphi\left( \cdot\right)$ is the neuronal input–output transfer function (applied element-wise), so $W \varphi(x)$ represents internally generated recurrent input.

Finally, $\eta(t)$ represents stochastic noise (e.g., Gaussian), capturing intrinsic fluctuations. Linearizing around a fixed point $x^{*}$ yields:

$\delta ẋ = J \delta x$	(2)

with Jacobian:

$J = \tau^{-1}\left[ -I + W diag\left( \varphi^{'}\left( x^{*} \right) \right) \right]$	(3)

Decomposition into eigenmodes ($\lambda_{k}$, $v_{k}$) gives modal relaxations:

$\delta x_{k\left( t \right)}\propto e^{\left\{ \lambda_{k}t \right\}}$	(4)

with characteristic timescales:

$\tau_{k}\approx\frac{-1}{Re\left( \lambda_{k} \right)}$	(5)

Fast modes correspond to eigenvalues with strongly negative real parts. Slow modes emerge when one or a few eigenvalues approach the boundary between stable and unstable dynamics, $Re\left( \lambda\right)\to0^{-}$, producing long correlation times and enhanced low‑frequency power, witnessed as Critical Slowing Down (CSD).This transition corresponds to the control parameter approaching its critical value (G → 1).

Connectivity statistics and structured (low‑rank) perturbations can reshape the spectrum, creating isolated spectral outliers that dominate long-timescale dynamics while most modes remain fast. This connection between slow recovery, rising variance/autocorrelation, and proximity to instability is central to quantitative CSD theory and is explicitly demonstrated for neural spiking transitions (Meisel et al., 2015). Random-matrix structure determines *which* slow modes are available. In large random networks with excitatory-inhibitory sign constraints and different column statistics, the eigenvalue cloud can deviate substantially from iid “circular-law” intuition, changing the density of decay rates and the likelihood of near-marginal (slow) directions. A second, complementary mechanism is that adding *structured* connectivity on top of randomness (often effectively low rank) can create spectral outliers: isolated eigenvalues separated from the bulk that dominate long-timescale dynamics while leaving most modes fast (Hu & Sompolinsky, 2022). In cortex, these ideas map naturally onto observations that population activity often rapidly collapses onto low-dimensional manifolds (a few dominant slow directions) while retaining fast fluctuations around them, consistent with effective low-dimensional dynamics in decision/ attention circuits. In cortical systems, these principles map naturally onto empirical findings that population activity often rapidly collapses onto low-dimensional manifolds (a few dominant slow directions), while retaining fast fluctuations around them. Finally, near-critical random recurrent networks provide a unified explanation for broadband field-potential spectra, in which slow activity amplification emerges as dynamical instability is approached. Thus, fast and slow modes are not separate mechanisms but rather distinct spectral components of the same recurrent architecture, determined by connectivity statistics, biophysical filtering, and proximity to dynamical instability, linking macroscopic “slow” power (Murray et al., 2014; Vidaurre et al., 2017), which is the primary focus of this study, to the same underlying eigenmode geometry.

**A two-mode Lorentzian approximation of random recurrent network PSDs**

The analytical two-mode Lorentzian approximation follows from linearizing the random recurrent network around a stable operating point. In that regime, the population dynamics can be written as a linear stochastic system, and after diagonalization of the effective connectivity matrix the activity decomposes into independent eigenmodes. Each mode relaxes exponentially with its own decay rate and, under approximately white stochastic drive, therefore contributes a Lorentzian term to the power spectral density (PSD). The full network spectrum is consequently a sum of modal Lorentzians, which can be approximated by a slow collective term and a fast bulk term when the eigenvalue spectrum is dominated by one near-critical slow mode together with a cloud of faster local modes.

$$ṙ(t) = A r(t) + \eta(t)$$

(6)

$$ẋₖ(t) = \lambdaₖ xₖ(t) + \etaₖ(t)$$

(7)

$$Sₖ(f) \propto1 / ((2\pi f)^{2} + \lambdaₖ^{2})$$

(8)

Accordingly, if the slow mode is sufficiently separated from the faster bulk, the PSD may be reduced to an effective two-mode form,

$$S(f) \approx C_{slow} / (f^{2}+ f_{slow}^{2}) + C_{fast} / (f^{2} + f_{fast}^{2})$$

(9)

This reduction is not exact, but rather a dominant-mode approximation that is expected to be most accurate when intermediate timescales are weak or clustered. Within this interpretation, the Lorentzian two-mode form provides a mathematically grounded reduced description of the broadband PSD shape of random recurrent networks near criticality. This account follows the analytical treatment of Chaudhuri et al. (2018) and is also consistent with later timescale-based frameworks using mixtures of Ornstein-Uhlenbeck processes, for which exponentially decaying modes likewise generate Lorentzian spectral components (Zeraati et al. 2021).

**No true common vertex is expected for isolated two-mode regional fits**

Although the two-mode Lorentzian model provides a useful compact description of regional PSDs, it does not in general support an exact common vertex when only $f_{slow,i}$ varies across regions. For two regions $i$ and $j$,
 $P_{i}\left( f \right)=A\left[ \frac{w}{f^{2}+f_{slow,i}^{2}}+\frac{1-w}{f^{2}+f_{\mathrm{fast}}^{2}} \right]$ and $P_{j}\left( f \right)=A\left[ \frac{w}{f^{2}+f_{slow,j}^{2}}+\frac{1-w}{f^{2}+f_{\mathrm{fast}}^{2}} \right]$. (10)

A shared vertex would require that all spectra intersect at the same frequency $f_{v}$, implying $P_{i}\left( f_{v} \right)=P_{j}\left( f_{v} \right)$ for all region pairs. Because the fast term is identical across regions, this condition reduces to

$\frac{1}{f_{v}^{2}+f_{slow,i}^{2}}=\frac{1}{f_{v}^{2}+f_{slow,j}^{2}}$ , (11)

which is satisfied only if $f_{slow,i}=f_{slow,j}$. Thus, once $f_{\mathrm{slow}}$ differs across regions, the analytical spectra generally cannot all pass through a single common vertex. At best, they may show approximate crossings over a restricted range, but not the exact shared intersection required for a true shape collapse.

This limitation is consistent with the interpretation of these modeled spectra as isolated regional, or “island,” approximations. In the Chaudhuri framework, the slow component is spatially distributed and becomes selectively enhanced when activity is averaged across a connected network, whereas isolated subnetworks do not preserve that same global averaging structure. Accordingly, the empirical cortex-wide PSDs may exhibit a much stronger approximate common-vertex organization than spectra generated from independently fitted local models.

Together, these results indicate that the analytical two-mode Lorentzian is useful as a reduced descriptive model of regional PSD shape, but it is not sufficient to generate a mathematically exact vertex-based collapse across regions when each spectrum is treated as an isolated unit with its own slow timescale. The residual non-collapse is therefore expected and supports the view that the empirical collapse reflects properties of the embedded large-scale cortical network rather than of independent local spectra alone.

**Figure S1. Influence of tSNR on the spatial organization of cortical fALFF.**
Group-averaged cortical maps of fALFF (A) computed from HCP resting-state data after exclusion of regions with low temporal signal-to-noise ratio (tSNR threshold set to 80). The spatial distributions closely match those obtained without tSNR filtering. (B) Distribution of fALFF peak values across individual subjects, indicating consistent rank-order of spectral distribution across participants (C) Split-half (odd vs. even) reliability estimates across subjects.

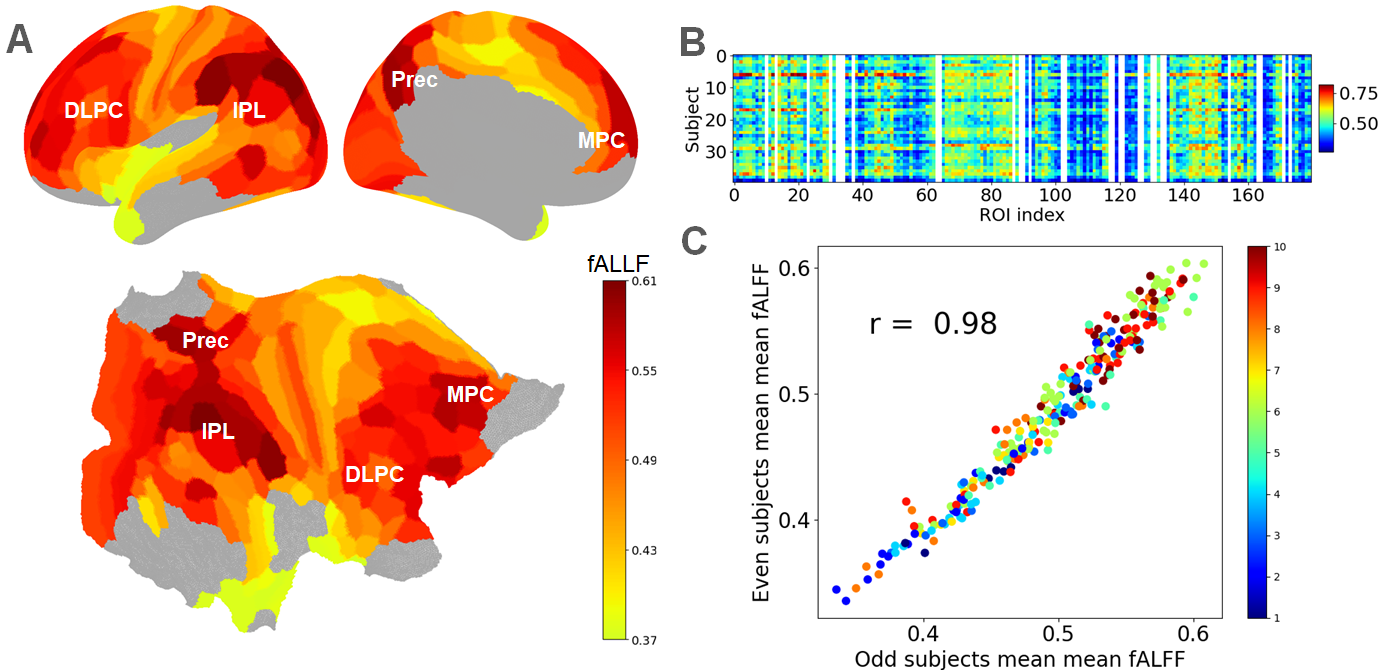

**Figure S2. Within-network consistency of regional fALFF distributions across subjects.**
Extending the pairwise subject-by-subject fALFF analysis shown in Figure 3C, we quantified the similarity of ROI-wise fALFF profiles within individual cortical networks. For each network, variance explained (R²) was computed across all subject-pair combinations using mean fALFF values across the network’s constituent ROIs. Bars summarize the mean paired R² values for all left-hemispheric ROIs (“All”) and for selected networks, including Auditory, Default Mode, Dorsal Attention, and Frontoparietal. Consistent with the whole-cortex analysis, regional fALFF organization remained broadly preserved across subjects, with the DMN showing particularly high consistency.

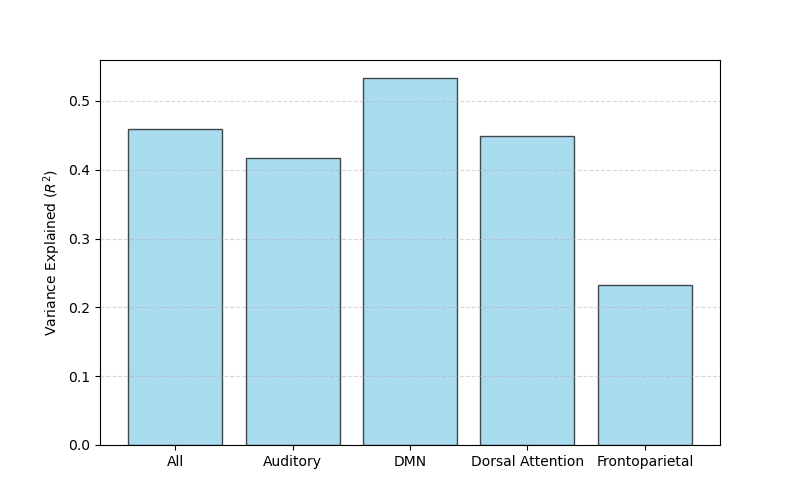

**Figure S3. Nonlinear dependence of simulated slow-timescale dynamics on proximity to criticality.** Simulations were performed while gradually increasing neural gain from **s**ub-critical to near-critical regimes. (A) Mean simulated fALFF as a function of the control parameter G (via gain), along with the fit equation. (B) Extending the same analysis across a broader gain range simulation, illustrates that nonlinear dependence is not confined to a narrow parameter interval.
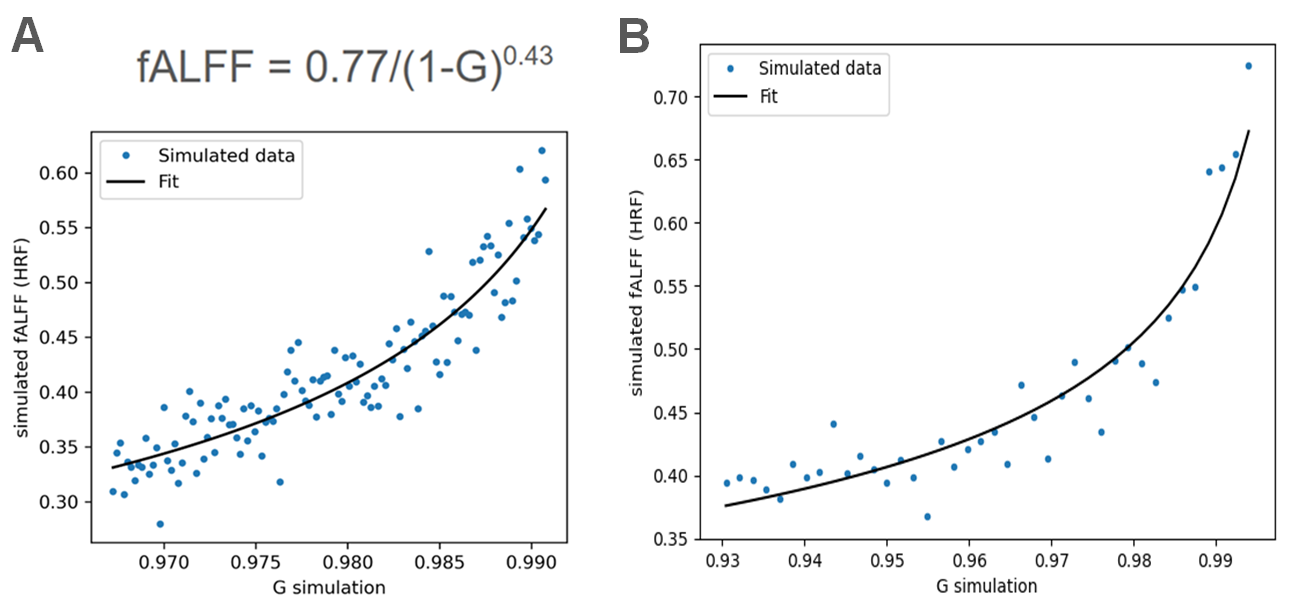

**Figure S4. Averaging across realizations improves simulation–data correspondence.**
Spatial correlation between simulated and empirical fALFF maps computed from a single realization (A), after averaging across three (B), and across five (C) independent realizations. Averaging reduces variance across simulations and increases correspondence with empirical cortical maps. The color coding depicts the control parameter G setting per each point in the simulated network. As can be seen there was an excellent fit between the simulation and HCP data. Of particular interest is the narrow range of the estimated G values: the vast majority of cortical dynamics could be well fitted within $\approx0.96-0.99$ , highlighting the strong sensitivity of the fluctuations to the network’s proximity to criticality.

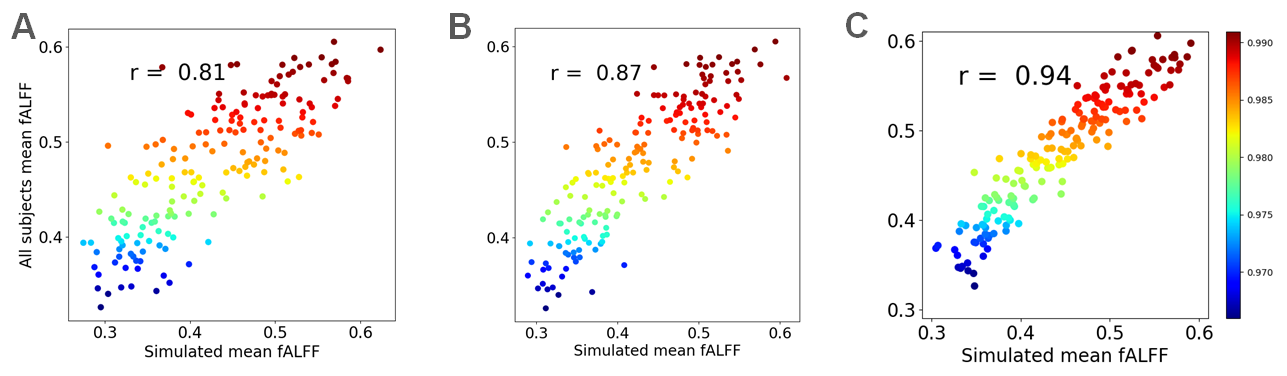

**Figure S5. Analytical approximation of the PSD profiles and their shape collapse.**

**(A)** Shape-collapse analysis of the analytical Lorentzian approximation, demonstrating partial alignment of regional PSD profiles when all model parameters were fit independently for each profile. **(B)** Gain-modulated analytical PSDs required additional adjustment of the Lorentzian amplitude to reproduce a common vertex point. **(C)** Even after amplitude adjustment, collapse of the analytical PSDs remained incomplete, indicating that simple modulation of reduced-model parameters is insufficient to capture the full organization of cortical spectral variation.

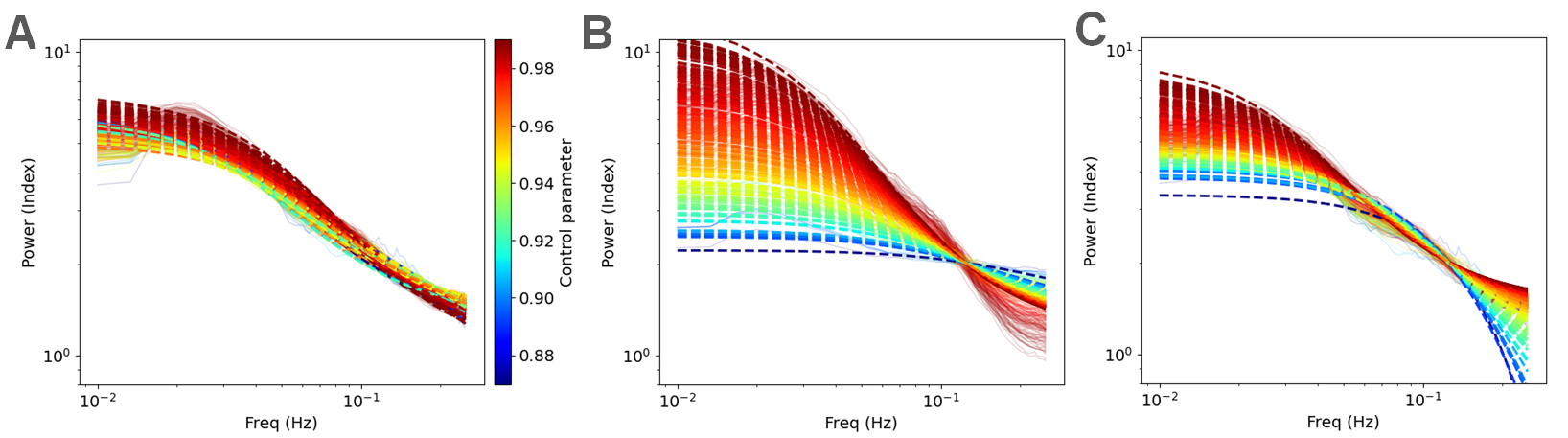

**Figure S6. Simulated recurrent networks approximate the structure of empirical cortical PSDs.**
**(A)** Pairwise spectral correlations across cortical ROIs, indicating strong similarity in spectral structure across regions despite differences in absolute fluctuation amplitudes. **(B)** Comparison of empirical PSD correlations with mean correlations across multiple realizations of simulated recurrent networks as a function of increasing control parameter G. **(C)** Simulated spectral profiles from isolated recurrent networks at different DTC, differing mainly in amplitude scale and lacking the common vertex observed in the underlying empirical cortical PSDs. Here, $\tau=60$ ms was chosen to better visualize the slow-frequency power fit, even though $\tau=20$ ms provided the best overall match across the per-gain spectral profiles.

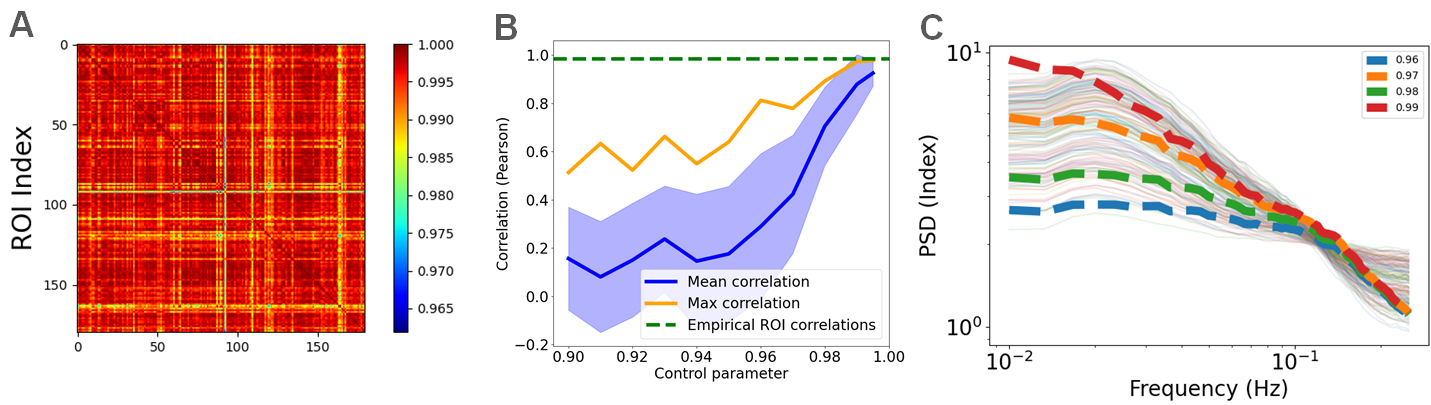

**Figure S7. Influence of tSNR on the spatial organization of cortical rFCD.**
Group-averaged cortical maps of rFCD (A) computed from HCP resting-state data after exclusion of regions with low temporal signal-to-noise ratio (tSNR thresholded to 80). The spatial distributions closely match those obtained without tSNR filtering. (B) Distribution of tSNR-corrected rFCD peak values across individual subjects, indicating consistent regional profiles of local functional connectivity across participants (C) Split-half reliability estimates across subjects.

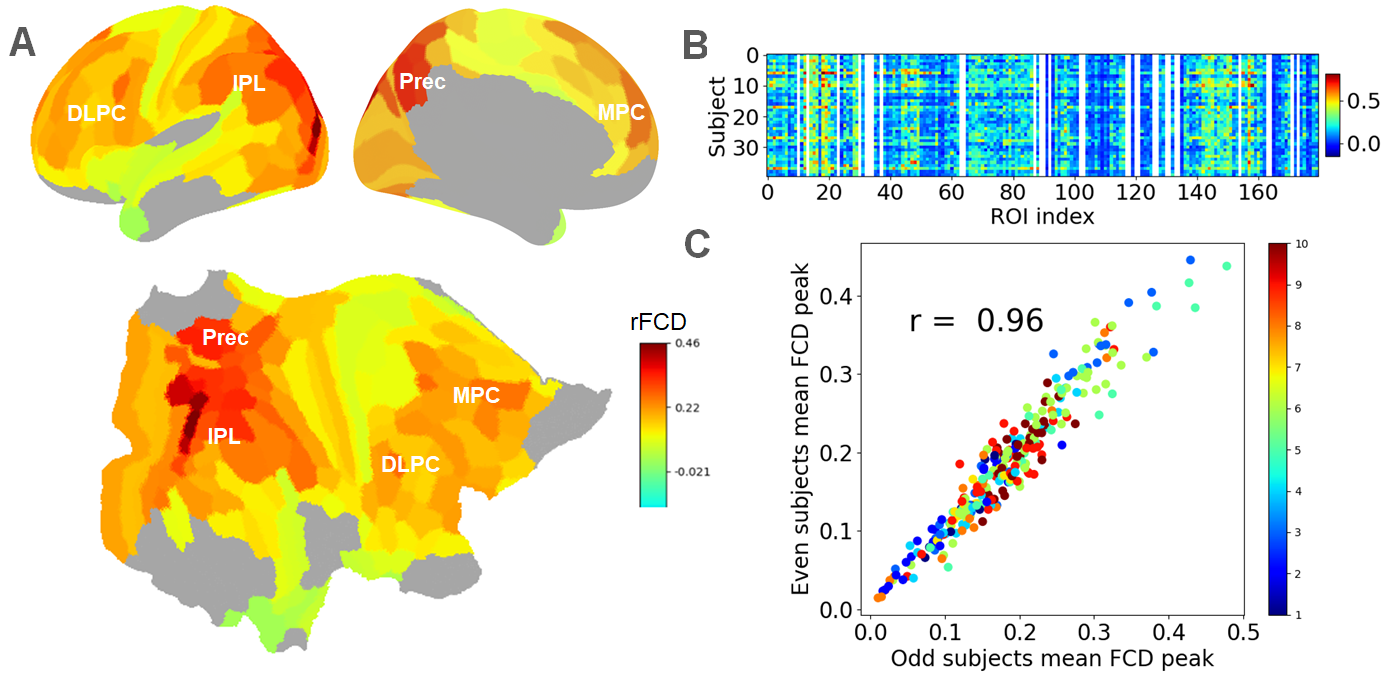

**Figure S8. Inter-subject consistency of fALFF–rFCD coupling**. fALFF–rFCD relationship across 40 subjects, indicating that the positive coupling between regional slow-timescale activity and local functional connectivity is consistently preserved across individuals, with Pearson correlation coefficients ranging from 0.70 to 0.94.

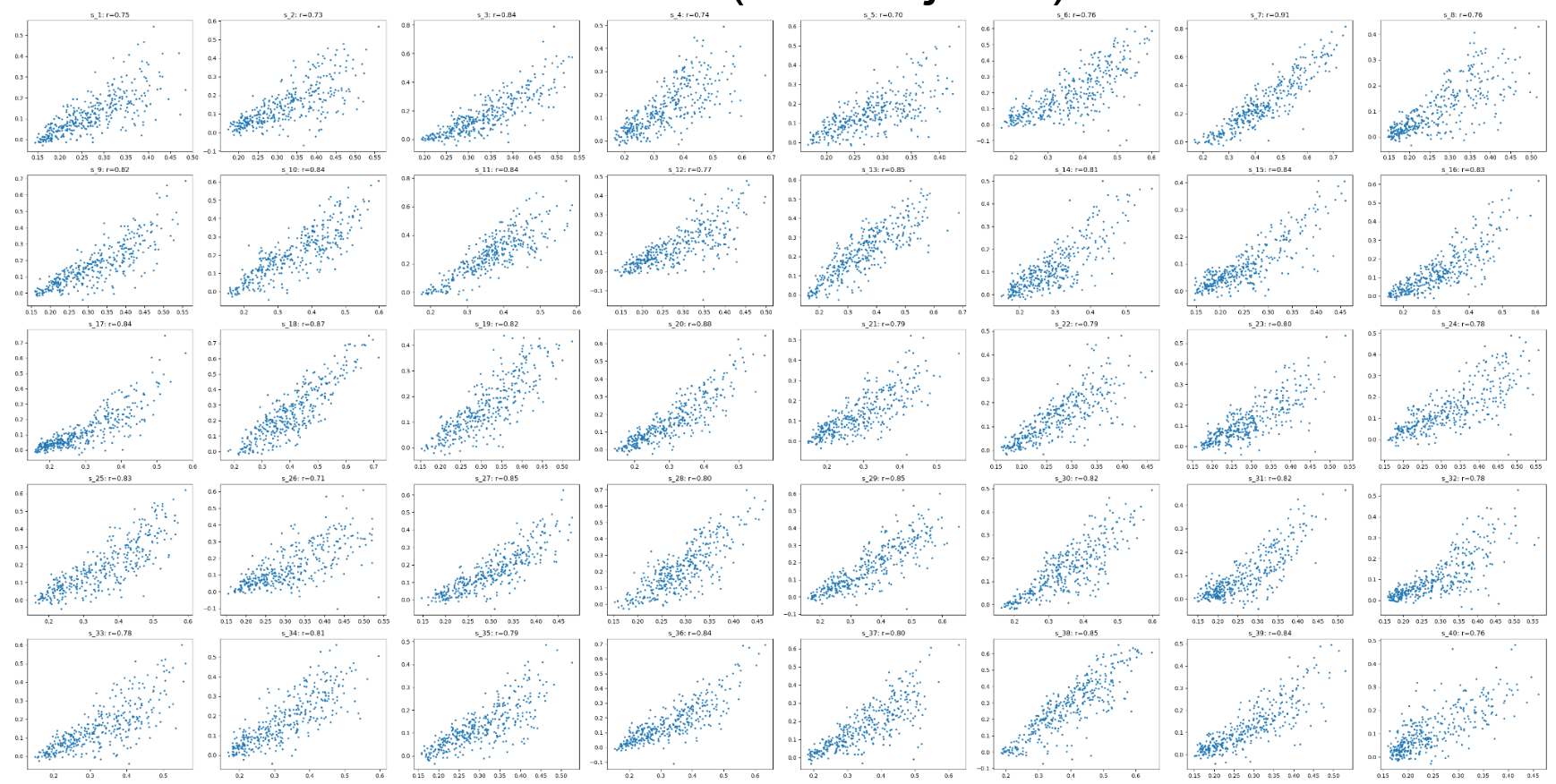
